# Fractalkine-induced microglial vasoregulation occurs within the retina and is altered early in diabetic retinopathy

**DOI:** 10.1101/2020.06.15.151464

**Authors:** Samuel A. Mills, Andrew I. Jobling, Michael A. Dixon, Bang V. Bui, Kirstan A. Vessey, Joanna A. Phipps, Ursula Greferath, Gene Venables, Vickie H.Y. Wong, Connie H.Y. Wong, Zheng He, Flora Hui, James C. Young, Josh Tonc, Elena Ivanova, Botir T. Sagdullaev, Erica L. Fletcher

## Abstract

Local blood flow control within the CNS is critical to proper function and is dependent on coordination between neurons, glia and blood vessels. Macroglia such as astrocytes and Müller cells, contribute to this neurovascular unit within the brain and retina, respectively. This study explored the role of microglia, the innate immune cell of the CNS, in retinal vasoregulation and highlights changes during early diabetes. Structurally, microglia were found to contact retinal capillaries and neuronal synapses. In the brain and retinal explants, the addition of fractalkine, the sole ligand for monocyte receptor Cx3cr1, resulted in capillary constriction at regions of microglial contact. This vascular regulation was dependent on microglial involvement, since mice lacking Cx3cr1, exhibited no fractalkine-induced constriction. Analysis of the microglial transcriptome identified several vasoactive genes, including angiotensinogen, a constituent of the renin-angiotensin system (RAS). Subsequent functional analysis showed that RAS blockade via candesartan, abolished microglial-induced capillary constriction. Microglial regulation was explored in a rat streptozotocin (STZ) model of diabetic retinopathy. Retinal blood flow was reduced after 4 weeks due to reduced capillary diameter and this was coincident with increased microglial association. Functional assessment showed loss of microglial-capillary response in STZ-treated animals and transcriptome analysis showed evidence of RAS pathway dysregulation in microglia. While candesartan treatment reversed capillary constriction in STZ-treated animals, blood flow remained decreased likely due to dilation of larger vessels. This work shows microglia actively participate in the neurovascular unit, with aberrant microglial-vascular function possibly contributing to the early vascular compromise during diabetic retinopathy.

**Significance Statement:** This work identifies a novel role for microglia, the innate immune cells of the CNS, in the local control of the retinal vasculature and identifies deficits early in diabetes. Microglia contact neurons and vasculature and express several vasoactive agents. Activation of microglial fractalkine-Cx3cr1 signalling leads to capillary constriction and blocking the renin-angiotensin system (RAS) with candesartan abolishes microglial-mediated vasoconstriction in the retina. In early diabetes, reduced retinal blood flow is coincident with capillary constriction, increased microglial-vessel association, loss of microglial-capillary regulation and altered microglial expression of the RAS pathway. While candesartan restores retinal capillary diameter early in diabetes, targeting of microglial-vascular regulation is required to prevent coincident dilation of large retinal vessels and reduced retinal blood flow.

## Introduction

The retina is one of the most metabolically active organs in the body, and in most mammals is supplied by an outer (choroidal) and inner (retinal) vascular network (1). While the choroid provides for the light-detecting photoreceptors within the outer retina, the retinal blood supply supports the numerous neurons and glia found in the ganglion cell and inner nuclear layers of the retina (2). The arterioles of the retinal blood supply enter at the optic disc and branch to form sequentially smaller vessels, including the retinal capillaries, establishing the superficial vascular plexus. These capillaries penetrate the inner retina, forming the relatively sparse intermediate vascular plexus, and deeper towards the outer retina forming the highly anastomosed deep vascular plexus. Completing the vascular circuit, blood returns via the venules on the retinal surface, which exit alongside the optic nerve (3, 4).

Blood flow throughout the retina is largely dependent on vessel calibre, which is tightly regulated to meet the metabolic demands of neuronal activity (5). An example of this is the well-defined hyperaemic response, whereby increased neuronal activity (via flickering light) results in arteriole dilation and increased inner retinal blood flow (6). Unlike peripheral blood vessels, retinal and brain vasculature have no direct neuronal input to modulate vascular tone, rather macroglial cells (Müller cells and astrocytes) are thought to actively regulate vascular calibre in response to changes in neural activity (7, 8). This type of coupling has given rise to the idea of a neurovascular unit, encompassing neurons, glia and blood vessels (7). While studies within the retina identified neuronal-dependent calcium increase in Müller cells to mediate vessel diameter change (9), more recent data suggest regulation of the inner retinal vasculature is more complex (10). Evidence for this comes from the fact that the same light stimulus can induce either vasoconstriction or vasodilation, and Müller cell-dependent calcium signalling only controls capillaries within the intermediate vascular plexus (11, 12). This suggests the existence of multiple regulatory pathways within the retina.

Recently it has been proposed that microglia, the innate immune cells of the retina, may also play a role in the neurovascular unit, although direct functional evidence is lacking (13). The conventional view of microglia is that they contribute to disease via the release of pro-inflammatory and neurotoxic cytokines (14–16). However, it is now recognised that microglia play several important, inflammation-independent roles in the normal brain and retina, such as dynamic synaptic surveillance and synaptic pruning (17–19). Despite this, the inflammation-independent response of microglia to neuronal signalling and their role in the regulation of vascular tone has yet to be confirmed.

While regulation of retinal blood flow is critical to retinal function (20), vascular dysfunction is known to occur in several pathologies, including diabetic retinopathy (DR). Early in the progression of DR, vascular pathology such as reduced retinal blood flow, micro-aneurysms and areas of vascular non-perfusion occur (21). Reduced retinal blood flow, in particular, presents early in humans with diabetes (22–24), and in animal models of diabetes (24). Altered inner retinal vascular regulation is considered a likely precursor to the development of severe vascular pathology in DR (25).

The present study investigates whether retinal microglia form a functional component of the neurovascular unit, and whether signalling through the fractalkine-Cx3cr1 pathway modulates vascular diameter. In addition, the work explores whether altered microglial involvement with the inner retinal vasculature may help explain the reduced retinal blood flow that occurs early during diabetes. Exploring the mechanisms responsible for the tight regulation between retinal neuronal activity and the local blood supply is critical to understanding retinal function in health and disease and may provide an empirical framework for future therapies targeting vascular pathogenesis.

## Results

### Microglia contact both retinal vasculature and neuronal synapses

Microglia within the CNS have a close association with the vasculature, particularly during injury and disease (26). However, less is known about microglial-vascular interactions in normal tissue. Within the retina, microglial cell bodies typically reside in the plexiform layers, while their processes extend throughout the retina (see *SI appendix,* Fig. S1). Inspection of the superficial vascular plexus shows microglia tiling the whole tissue (Fig. 1*A*, Cx3cr1^GFP/+^ mouse retina, EGFP, green) and in close association with retinal vasculature (Fig. 1*A* *inset*; IB4, red). When microglial process contact with retinal vessels of different diameters is quantified relative to the respective area of each vessel diameter class, microglia are seen to interact with smaller retinal vessels (≤15μm), particularly the smallest retinal capillaries (<10μm), when compared to the larger vessels (Fig. 1*B*; one-way ANOVA, p < 0.05, 0.001 for 15-20μm and >20μm, respectively). At the ultrastructural level (Fig. 1*C*; Cx3cr1^GFP/+^ mouse retina), a microglial process (MC, stained for EGFP) abuts a pericyte (PC), which lies over an endothelial cell (EC) lining the capillary lumen (CL). This microglial-pericyte contact is also investigated immunohistochemically using the NG2-DsRed reporter mouse, which labels pericyte somata and processes (Fig. 1*D*, red). A microglial cell (Iba-1, green) is observed to make contact with two pericyte somata (red), with nuclei immunolabelled with DAPI (blue). Orthogonal projections (top and right) from the boxed area, show direct contact between the two cell types. The contact indicated with the asterisk was further imaged at higher resolution to show direct contact between the microglial process (green) and the pericyte soma (red; asterisk in Fig. 1*E*, also see *SI appendix,* Fig. S2 and Video S2). The extent of microglial contact with pericytes somata, processes (NG2-labelled) and capillary areas devoid of pericyte contact (NG2 negative / IB4 positive regions) was quantified in rat retina, with no preference observed for microglial-pericyte or microglial-vessel contact (Fig. 1*F*). In addition to contacting retinal vessels (IB4, magenta, asterisk in Fig. 1*G*), microglia (EGFP, green) are also observed to extend processes into the inner plexiform layer (IPL), where neuronal synapses reside (Fig. 1*G*, VGLUT1 red, arrow heads; DAPI blue). The *inset* shows a rendering of these microglial-neuronal interactions at higher magnification. This is also observed in the human retina (Fig.1*H*, DAPI, blue) with microglia (Iba-1, green) contacting both retinal vessels (vitronectin, magenta, asterisk) and neuronal synapses (VGLUT1, red. arrow heads). When quantified in the *Cx3cr1^+/GFP^* mouse retinae, the majority of microglia (EGFP, green) in the inner retina contact both neuronal synapses (VGLUT1, blue) and retinal vessels (IB4, red; Fig. 1*I* inset, 73 ± 13%, rat retina). All individual channels for immunolocalization are shown in *SI appendix* (Fig. S3)

**Figure 1.**
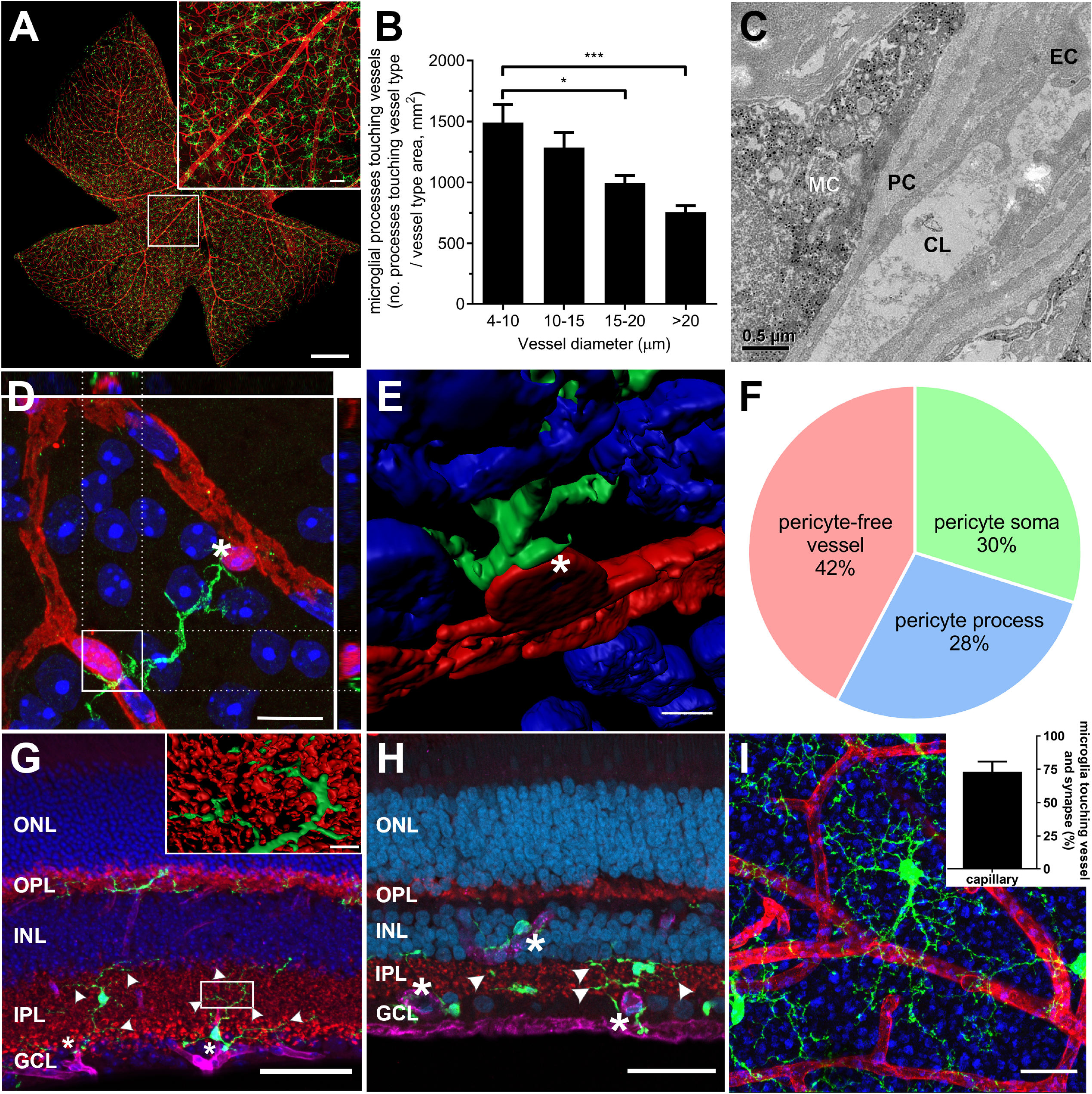
Retinal microglia associate with vasculature and neuronal synapses. *A:* Wholemounted mouse retina (*Cx3cr1^GFP/+^*) was labelled with anti-EGFP (microglia, green), and *Griffonia simplicifolia* isolectin B4 (IB4, blood vessels, red). The highlighted region is magnified showing microglial association with vessels within the superficial vascular plexus (*inset*). *B:* The association of microglial processes with vessels of different diameters within the superficial plexus was quantified relative to vessel area for each vessel size and show microglia preferentially associate with capillaries. *C:* The ultrastructure of microglia-vessel contact within the *Cx3cr1^GFP/+^* retina shows microglial processes (immunolabelled against EGFP, black dots) adjoin pericytes, which contact the endothelial cells lining the capillary lumen. *D:* A wholemounted retina from the NG2-DsRed pericyte reporter mouse (pericyte somata, processes, red) stained with Iba-1 (microglia, green) and DAPI (nuclei, blue) shows a microglial process making contact with pericyte somata. The boxed region is shown in XZ and YZ orthogonal projections (above and right). *E*: A high resolution rendered image of microglial-pericyte contact taken from asterisk in panel *D*. *F*: Microglial-pericyte interaction was further probed in rat retina and the extent of contact with pericyte somata, processes (NG2 +ve) and capillary areas lacking pericyte contact (NG2 −ve / IB4 +ve) quantified. *G:* A vertical section from a *Cx3cr1^GFP/+^* retina labelled for blood vessels (IB4, magenta), microglia (EGFP, green) neuronal synapses (VGLUT1, red) and cell nuclei (DAPI, blue), showing microglial processes contact retinal vessels (asterisk) and neuronal synapses (arrow heads). The boxed region was imaged at higher resolution and rendered to highlight microglial-synapse interaction (*inset*). *H*; Neuronal-microglial-vascular contact is also observed in human retina (microglia, Iba-1, green; vessels, vitronectin, magenta, asterisk; neuronal synapses, VGLUT1, red, arrow heads; cell nuclei, DAPI, blue). *I:* When neuronal-microglial contact was quantified in the *Cx3cr1^+/GFP^* mouse at the level of the inner retina (vessels, IB4, red; microglia, EGFP, green, VGLUT1, blue), the majority of microglia contact both neuronal synapses and retinal vessels. Data presented as mean ± SEM, n= 5 (*B, F*), n=3 (*I inset*), *p<0.05, ***p<0.001. MC, microglia; PC, pericyte; EC, endothelial cell; CL, capillary lumen; ONL, outer nuclear layer; OPL, outer plexiform layer; INL, inner nuclear layer; IPL, inner plexiform layer; GCL, ganglion cell layer. Scale bars 500μm (*A*), 50μm (*A inset, G*, *H), 20*μm *(I*), 10μm (*D*), 5μm (*E*) 0.5μm (*C*).

### Microglia modulate vessel diameter and express vasoactive genes

Within the brain and retina, macroglial (astrocyte and Müller cell) cell contact with neuronal synapses and vasculature is critical for local control of blood supply in response to neuronal activity (7, 8). To determine whether microglia play a similar role, *Cx3cr1^GFP/+^* retinae were isolated and maintained e*x vivo*. Microglia were visualised via their expression of EGFP (Fig. 2*A*; green) and vessels were labelled with rhodamine B (Fig. 2*A*; red). As the fractalkine-Cx3cr1 axis is thought to mediate neuronal-microglial communication, blood vessels and microglia were imaged while fractalkine (200ng/ml) or PBS was perfused into the chamber (*SI appendix,* Video S1). Vessel diameter change was monitored and expressed relative to the baseline value for the same region of vessel.

**Figure 2.**
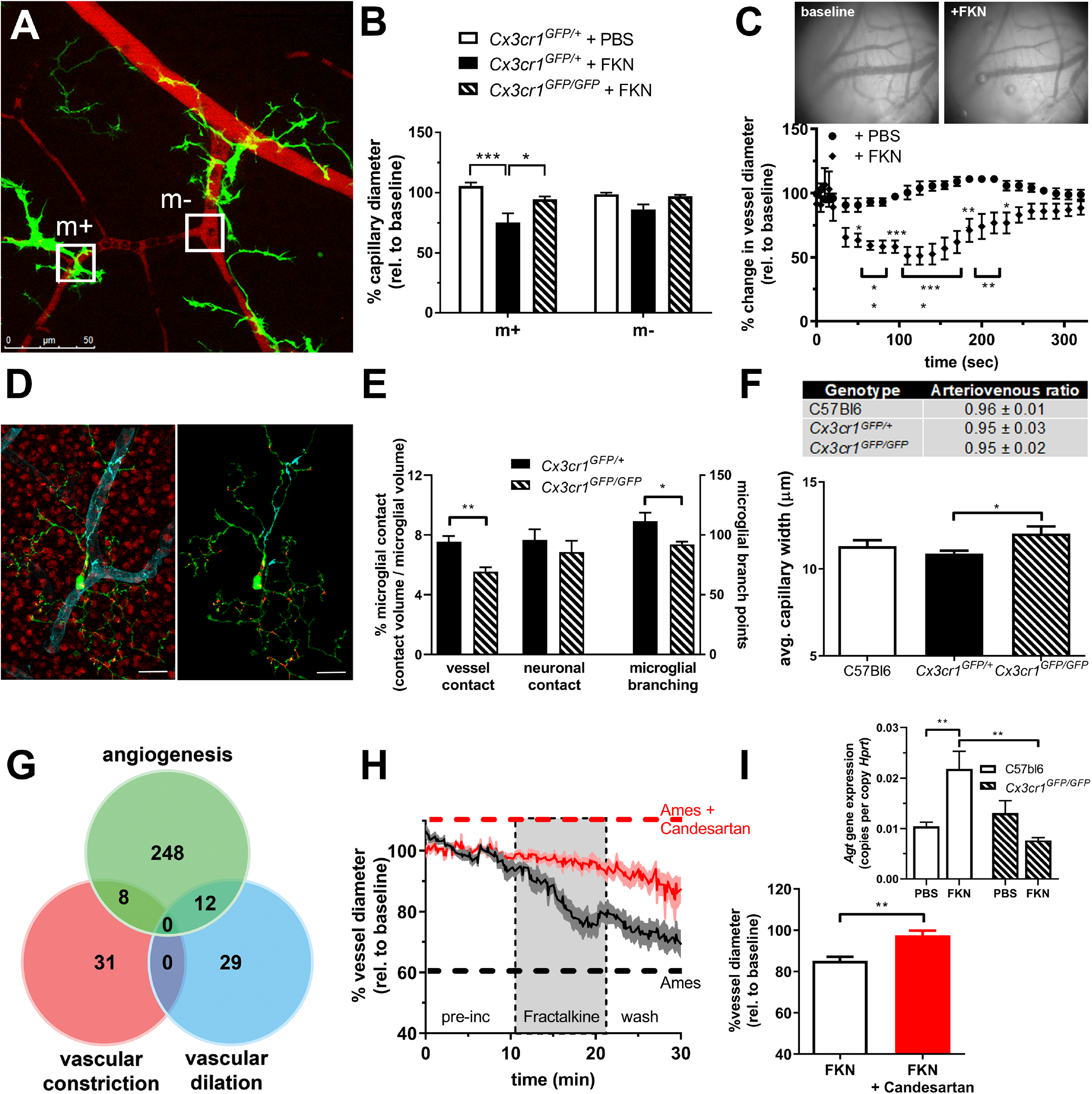
Microglia constrict retinal capillaries via fractalkine-Cx3cr1 signalling and express genes for vasoactive agents. *A: Ex vivo Cx3cr1^GFP/+^* retinae (EGFP; microglia, green) were labelled with Rhodamine B (blood vessels, red) and imaged under live cell microscopy. *B:* The addition of fractalkine (FKN, 200ng/ml) induced vasoconstriction at sites of microglial contact (m+. *n* = 4 PBS, *n* = 6 FKN), while no significant vessel alteration occurred in areas lacking microglial processes (m−, *n* = 5 PBS, n=6 FKN). When performed on *Cx3cr1^GFP/GFP^* retinae, no constriction was evident (*n* = 5). *C:* The response of brain vasculature to fractalkine was tested in rat thin scull preparations, with constriction evident 120 seconds post-injection (*n* = 3 PBS, FKN). The insets show representative images at baseline and after fractalkine addition. *D:* Retinal microglia (EGFP, green), neuronal synapses (VGLUT1, red) and blood vessels (IB4, light blue) were imaged in *Cx3cr1^GFP/+^* and *Cx3cr1^GFP/GFP^* animals and the extent of vascular and neuronal contact quantified relative to microglial volume (see isolated microglia, red - neuronal contacts; blue - vascular contacts). *E:* Grouped data showed *Cx3cr1^GFP/GFP^* retinae to have reduced vascular contacts compared to *Cx3cr1^GFP/+^* retinae (*n*=5), while there was no difference in neuronal contacts. *Cx3cr1^GFP/GFP^* microglia exhibited reduced process branching (*n*=5). *F:* Using *in vivo* OCTA, retinal capillary diameter was increased in *Cx3cr1^GFP/GFP^* animals compared to *Cx3cr1^GFP/+^* retinae (*n*=4 C57Bl6, *n*=6 *Cx3cr1^GFP/+^*, *Cx3cr1^GFP/GFP^*), while there was no alteration in the diameter of arterioles or venules (A/V ratio shown in table, *n*=4 C57Bl6, *n*=6 *Cx3cr1^GFP/+^*, *n*=5 *Cx3cr1^GFP/GFP^*). *G:* RNAseq was performed on FACS-isolated rat retinal microglia, with 268 genes identified as being angiogenic (GO:0001525), while 39 genes were involved in vascular constriction and 41 genes in vascular dilation (regulation of blood vessel diameter, GO:0097746). *H*: Vessel diameter was quantified in rat retinal explants pre-incubated in Ames (black trace) or Ames + 230 nM candesartan (red trace) for 10 minutes, after which fractalkine (FKN, 200 ng/ml) was added (shaded area, representative data from 1 retina, *n* = 5 vessels). *I*: When grouped data were analysed 10 minutes after fractalkine addition, constriction was abolished when pre-incubated with candesartan (*n* = 7 fractalkine, *n* = 5 fractalkine + candesartan). Further supporting a role for the RAS, *ex vivo* incubation with fractalkine (FKN) resulted in an increase in microglial *Agt* expression, while this was not evident in the microglia isolated from *Cx3cr1^GFP/GFP^* retinae (*inset*, *n*=6). Data expressed as mean ± SEM, \**p<*0.05, **p<0.01, ***p<0.001, ****p<0.0001. Scale bar 50μm (*A*), 15μm (*D*).

In response to fractalkine, blood vessel regions that were associated with microglial processes (m+) constricted (Fig. 2*B* m+; 2-way ANOVA; PBS versus fractalkine, *p* < 0.001), while those regions that were further away from microglial processes (m−) exhibited no significant alteration in capillary diameter (Fig. 2*B* m−; 2-way ANOVA; PBS versus fractalkine, *p* = 0.26). These *ex vivo* preparations showed minimal microglial process movement at the vascular level throughout the imaging, including during fractalkine exposure (*SI appendix,* Video S1 and Fig. S4). When explants taken from animals lacking Cx3cr1 (*Cx3cr1^GFP/GFP^*) were exposed to fractalkine, no alteration in vessel diameter was observed compared to PBS controls at regions with (m+; 105.7± 2.7% versus 94.7 ± 2.3%, 2-way ANOVA p=0.52) or without (m−; 98.8 ± 1.2% versus 97 ± 1.3%, 2-way ANOVA p=0.999) microglial contact (Fig. 2*B*). Finally, to explore whether this vasomodulatory function of fractalkine was retina-specific, superficial vessels within the rat brain were imaged using a thin skull preparation. These preliminary data showed that while vehicle delivery resulted in no alteration in vessel diameter, the subdural addition of fractalkine lead to a significant constriction of the smaller vessels (Fig. 2 *C*; RM 2-way ANOVA, vessels ≤ 15μm, p < 0.05). While both tissues show a fractalkine-induced constriction, the difference in vessel kinetic response likely reflects the different systems used to explore microglial vasoregulation (*ex vivo* and *in vivo,* respectively).

Since the *Cx3cr1^GFP/GFP^* retina showed no fractalkine-induced vessel constriction, microglial contact with retinal vessels and neurons was explored. High resolution immunocytochemical analysis of microglia (EGFP, green) contact with neuronal synapses (VGLUT1, red) and vessels (IB4, light blue) was undertaken to enable specific areas of contact to be quantified (Fig. 2*D*). When the volume of contact per individual microglia was calculated, *Cx3cr1^GFP/GFP^* animals had fewer vessel contacts than animals with one functional copy of Cx3cr1 (Fig. 2*E*; *Cx3cr1^GFP/+^* 7.5 ± 0.4% versus *Cx3cr1^GFP/GFP^* 5.5 ± 0.3%, *t*-test p=0.004). While there was no difference in neuronal contacts between the two genotypes, *Cx3cr1^GFP/GFP^* animals showed less microglial process branching (Fig. 2*E*; *Cx3cr1^GFP/+^* 111.5 ± 7.2 versus *Cx3cr1^GFP/GFP^* 92.2 ± 2.1, *t*-test p=0.03), reflecting the literature showing *Cx3cr1^GFP/GFP^* to have a more activated inflammatory profile (28). When retinal capillary diameters were compared to C57bl6 control animals, *Cx3cr1^GFP/+^* capillaries were similar to controls (Fig. 2*F*; C57bl6 11.3 ± 0.3μm versus *Cx3cr1^GFP/+^* 10.9 ± 0.2μm, 1-way ANOVA p=0.66), while *Cx3cr1^GFP/GFP^* showed increased capillary diameters (Fig. 2*F*; *Cx3cr1^GFP/+^* 10.9 ± 0.2μm versus *Cx3cr1^GFP/GFP^* 12 ± 0.4μm, 1-way ANOVA p=0.047). There was no difference in larger vessel diameter for any genotype (Fig. 2*F* *inset*; p=0.87 and 0.94 for *Cx3cr1^GFP/+^* and *Cx3cr1^GFP/GFP^,* respectively).

RNA-Seq was performed on FACS-isolated microglia collected from 12-week-old dark agouti rats to determine whether vasomodulatory factors were contained within the microglial transcriptome. To confirm the purity of sample, the mapped genes were compared to a published list of microglial markers (29), with 23/29 markers identified in our gene population, including the microglial-specific marker *Tmem119* (*SI appendix,* Table S1)(41). The microglial transcriptome was also compared to microglial-enriched genes reported in several studies, with significant overlap observed, while there was little contamination from known neuronal genes (*SI appendix,* Fig. S5). The expressed gene population was compared against genes known to be involved in angiogenesis (GO:0001525, 407 genes) and regulation of blood vessel diameter (GO:0097746, 310 genes). In total, 268 genes expressed in the microglial population were identified to have roles in angiogenic pathways (Fig. 2*G*, and *SI appendix*, table S2), such as hypoxia inducible factor 1 alpha (*Hif1a)* and vascular endothelial growth factor A and B (*Vegf A/B)*. When vessel diameter regulation was explored, 41 genes were found to have a role in vasodilation such as phospholipase A2 (*Pla2g6*) and sirtuin 1 (*Sirt1*), while 39 genes were identified with vasoconstriction, including endothelin 1, 3 (*Edn1, 3*) and arachidonate 5-lipoxygenase (*Alox5*) and angiotensinogen (*Agt*; Fig. 2*G*, and *SI appendix*, tables S3 and S4, respectively).

As angiotensinogen is a constituent of the renin-angiotensin system (RAS), which is involved in retinal vessel regulation via the angiotensin II receptor type 1 (AT1R) (30, 31), *ex vivo* experiments were performed using the AT1R antagonist, candesartan. Baseline capillary diameter was averaged over 10 minutes in rat retinal explants exposed to Ames (black trace) and Ames + candesartan (230 nM; red trace) and after which time fractalkine was added (shaded area in Fig. 2*H*). Similar to that observed in the *Cx3cr1^GFP/+^* mouse (Fig. 2*A*, *2B*), exposure of the rat retinae to fractalkine induced capillary constriction, while exposure to candesartan blocked any fractalkine-induced constriction (Fig. 2*H*). When grouped data were analysed, candesartan abolished the fractalkine-induced vasoconstriction (Fig. 2*I*, *t*-test, p<0.01). To further support the role of RAS in microglial-mediated vessel regulation, control C57bl6 and *Cx3cr1^GFP/GFP^* were exposed *ex vivo* to fractalkine (FKN) for 2 hours, microglia isolated and the expression of angiotensinogen (*Agt*) quantified (Fig. 2*I* *inset*). While exposure to fractalkine increased *Agt* expression in control retinae, *Cx3cr1^GFP/GFP^* retinae which previously exhibited no microglial-mediated constriction (Fig. 2*B*), showed no expression change (Fig. 2*I* *inset*; +FKN, C57bl6 21.8 ± 3.5 copies/1000 copies *Hprt* versus *Cx3cr1^GFP/+^* 7.7 ± 0.6 copies/1000 copies *Hprt*, 2-way ANOVA p=0.017). The current data show that microglia are capable of modulating vascular constriction within the retina and broader regions of the CNS via the fractalkine-Cx3cr1 pathway. While they express several gene transcripts for known vasoactive agents, microglial regulation of retinal vessels occurs via AT1R activation.

### Retinal blood flow and capillary diameter is changed in early diabetes

The regulation of retinal blood supply is critical to normal function, with retinal pathologies, such as DR, exhibiting early retinal blood flow defects and abnormal neurovascular coupling (22, 24, 32). To explore whether microglial vasoregulation was altered during early diabetes, adult dark agouti rats were rendered diabetic via a single injection of STZ with significant hyperglycaemia evident throughout the 4-week experimental period (*SI appendix*, table S5).

As reduced retinal blood flow is a consistent and early alteration in patients with diabetes and animal models (23, 24), quantitative vessel-dependent kinetic analysis using sodium fluorescein (33) was used to confirm vascular dysfunction. Average normalised fluorescence intensity was calculated over time for every pixel within the fundus image (see Fig. 3*A, B, C* *insets*), grouped on vessel type, and *en face* heat-maps produced (Fig. 3*A*, *B*, *C*, fill times), with warmer colours indicating greater time taken to fill (slower blood flow). Vessel-dependent kinetic analysis revealed arterioles in STZ-treated animals took longer to fill (Fig. 3*D*; median regression analysis, *p* < 0.05), reflecting reduced blood flow. Due to the serial nature of the retinal vasculature, this increase in fill time was also observed in retinal capillaries and venules (Fig. 3*D*; median regression analysis, *p* < 0.05), with no vessel-specific deficit identified (median regression analysis, *p* > 0.05). Drain times were also longer in all retinal vessels (Fig. 3*E*; median regression analysis, *p* < 0.05), with the effect significantly greater than that observed for fill times (median regression analysis, *p* < 0.05). The reduced arteriolar and venular blood flow in STZ-treated animals was verified using velocimetry (*SI appendix,* Fig. S6) and the clinically relevant arterio-venous transit time was also exhibited reduced blood flow (increased transit time, *SI appendix*, Fig. S6*D*). The decrease in retinal blood flow kinetics was independent of systemic change, with systolic blood pressure, blood haematocrit and intraocular pressure unaltered (*SI appendix,* Fig. S7).

**Figure 3.**
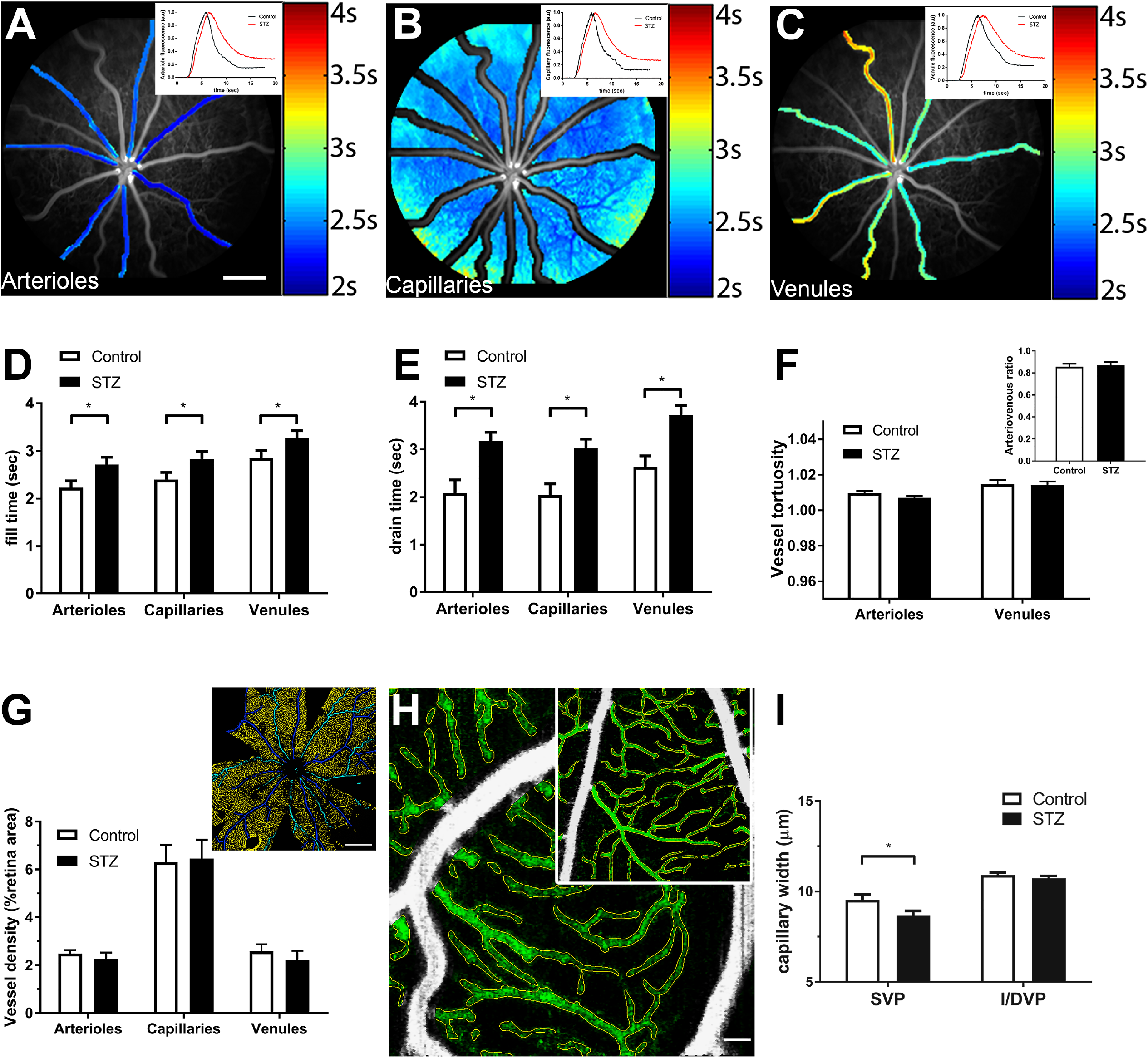
Retinal blood flow is reduced and capillaries are constricted after 4 weeks of diabetes. VFA was used to quantify retinal blood flow in control and STZ-treated animals. *A-C*: En face heat-maps depicting fill time for arterioles, capillaries and venules, with insets showing representative average normalised fluorescence intensity traces for control (black line) and STZ-treated (red line) animals. *D, E*: The times taken to reach half maximum intensity (*D*, fill time) and half of final value from maximum (*E*, drain time) were quantified, showing fill and drain times were significantly increased in all vessel types in STZ-treated animals (unfilled bars, control *n* = 23; filled bars STZ, *n* = 21). *F*: Sodium fluorescein fundus images were quantified for large vessel tortuosity (*n* = 13) and arteriovenous ratio (*inset*, *n* = 13), with no difference observed between STZ-treated (filled bars) and control (unfilled bars) animals. *G:* Immunohistochemistry was used to quantify vascular density in control and STZ-treated (unfilled and filled bars) eyes, with no difference observed between the two groups (*n* = 11). The rendered image shows the segmented vessel types (*inset*, capillaries in yellow, arterioles in blue and venules in cyan). *H:* OCTA was performed *in vivo* to measure capillary diameter in control and STZ-treated (*inset*) animals, with the vessels measured shown in green. *I*: A significant decrease was observed in the capillary diameter in STZ-treated animals (*n* = 12) compared to control (*n* = 10) within the superior vascular plexus. No alteration was observed in the intermediate/deep vascular plexi. Group data expressed as mean ± SEM. *p<0.05. Scale bars 500 μm (*A*), 1 mm (*G*), 50 μm (*H*).

As vessel change affects blood flow in DR (34, 35), the morphology of large diameter vessels was assessed from fluorescein images at peak fluorescent intensity. No change in retinal large vessel tortuosity (Fig. 3*F*; 2-way ANOVA, arterioles *p* = 0.52, venules *p* = 0.98), or arteriole / venule diameter (arteriovenous ratio, Fig. 3*F* *inset*; *t*-test, *p* = 0.48) was observed between the two cohorts of animals. Similarly, when arteriole, capillary and venule densities were separately quantified using retinal wholemount immunohistochemistry (Fig. 3*G* *inset* shows the rendered image of arterioles, dark blue; venules, cyan; and capillaries, yellow), no change in vessel densities were observed between control and STZ-treated animals (Fig. 3*G*, 2-way ANOVA, arterioles *p* = 0.98, venules *p* = 0.99, capillaries *p* = 0.94). As fluorescein image analysis and immunohistochemistry lack the resolution to assess capillary diameter, OCTA was used to quantify this *in vivo*. Images of the superficial retinal capillary network were obtained for control (Fig. 3*H*) and STZ-treated (Fig. 3*H* *inset*) animals and quantification (green overlay showing measured capillaries) revealed a decrease in capillary diameter in the STZ-treated cohort (Fig. 3*I*; 2-way ANOVA, *p* < 0.05). When a similar analysis was performed on the intermediate and deep capillary plexi (I/DVP), no alteration in diameter was detected (Fig. 3*I*; 2-way ANOVA, p=0.72).

In summary, retinal blood flow was significantly slower in diabetes, with *in vivo* OCTA revealing retinal capillary constriction within the superficial vascular plexus 4 weeks after STZ-induced diabetes. These diameter changes were restricted to the capillary network, as larger vessels remained unaltered and there was no change in retinal vascular coverage.

### Retinal microglia contact with capillaries and pericytes is increased in early diabetes, independent of activation

The extent of microglial (Fig. 4*A* *inset*, green, Iba-1) contact with arterioles, capillaries and venules (Fig. 4*A* *inset* red, IB4) was quantified for control and STZ-treated animals to determine whether the retinal capillary constriction in diabetes was accompanied by altered microglial association. While microglia exhibited a similar association with large diameter arterioles and venules (Fig. 4*A*; 2-way ANOVA, *p* > 0.99 and *p* > 0.66, respectively), microglial-capillary association was increased in STZ-treated animals (Fig.4*A*; 2-way ANOVA, *p* < 0.05). In addition, microglial-pericyte association (Fig. 4*B* *inset* microglia green, Iba-1; pericytes light blue, NG2, vessels red, IB4) was increased within the central retina of STZ-treated animals (Fig. 4*B*, 2-way ANOVA, p<0.05). There was no vessel dropout (Fig. 3*G*), nor loss of retinal pericytes (*SI appendix,* Fig. S8) at this early stage of diabetes. The association of microglia with pericytes and capillary areas lacking pericyte contact was further explored in control and STZ-treated animals using quantitative image analysis (Fig. 4*C* *inset,* rendered image showing pericyte somata red; pericyte processes green; pericyte-free vessel blue and skeletonised microglia). While quantitative analysis showed no specific preference for microglia to contact pericyte somata, processes or capillary areas lacking pericytes (Fig.4*C*; 2-way ANOVA, *p* = 0.16), there was increased microglial association with all three at 4 weeks of diabetes (Fig.4*C*; 2-way ANOVA, *p* < 0.01). To determine whether this microglial effect was specific, or a result of a more generalised macroglial response as has been shown in later stages of diabetes (36, 37), astrocyte density and Müller cell gliosis were quantified. Vessel-specific astrocyte coverage (Fig 4*D*) and Müller cell gliosis (Fig. 4*E*) were unaltered after 4 weeks STZ treatment (2-way ANOVA, *p* > 0.92 and 0.99 respectively).

**Figure 4.**
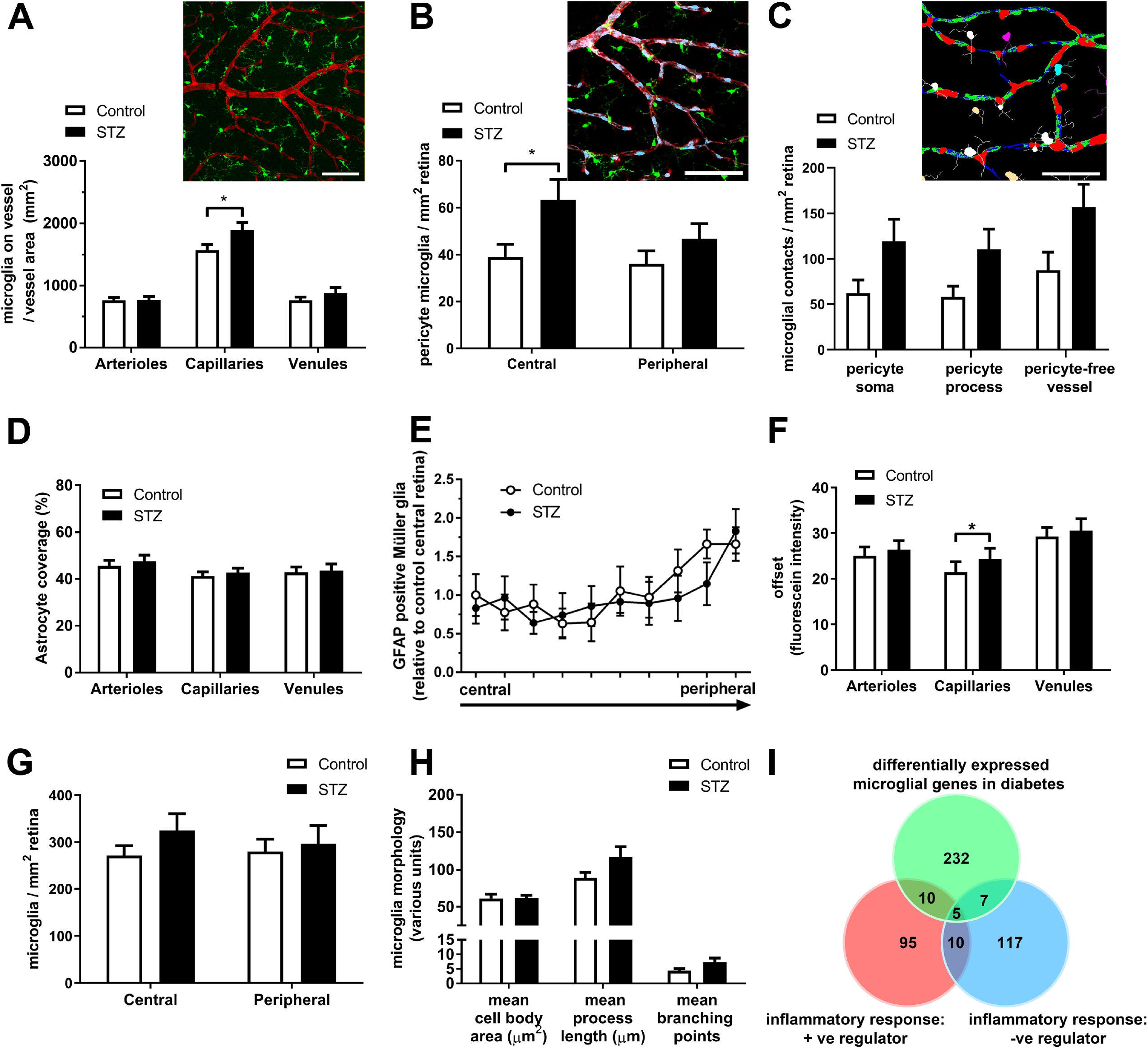
Microglia increase their contact with retinal capillaries after 4 weeks of STZ-induced diabetes. *A*: Wholemounted retina from control (*inset*) and STZ-treated animals were labelled for Iba-1 (microglia, green) and IB4-FITC (blood vessels, red) and the extent of contact between microglia and vasculature quantified for each vessel type. While no difference in large vessel contacts occurred, microglia-capillary contact increased in the central retina of the STZ-treated animals (filled bars, *n* = 11). *B:* Control (*inset*) and STZ-treated animals were labelled for Iba-1 (microglia, green), NG2 (pericytes, light blue) and IB4-FITC (blood vessels, red) and the extent of microglia-pericyte contact quantified for each vessel type. Microglial-pericyte association increased within the central retina of STZ-treated animals (filled bars, *n* = 11). *C*: Using similar immunolabelling as in *B,* microglial association with pericyte somata, processes and capillary areas lacking pericyte contact was quantified. The image analysis render (*inset*) highlights pericyte somata (red), pericyte processes (green) and pericyte-free vessels (blue), while microglia touching each of these regions were skeletonised and colour coded for quantification. While there were no preferential association, all contacts were increased in STZ-treated (filled bars, *n* = 5) compared to control (unfilled bars, *n* = 5) retinae. *D, E*: Macroglial change was assessed in control (unfilled bars, *n* = 11) and STZ-treated (filled bars, *n* = 11) retinae, with no alteration in astrocyte coverage (*D*), nor Müller cell gliosis (*E*) observed (*n* = 6). *F*: Kinetic analysis of VFA was used to quantify fluorescein offset as a measure of BRB integrity. While arterioles and venules showed no change, capillary offset was increased in STZ-treated animals (unfilled bars, control *n* = 23; filled bars STZ, *n* = 21). *G, H:* The inflammatory status of microglia was assessed morphologically and no difference was found in the number of monocytes / microglia in central and peripheral retina (*G*, *n* = 11), cell soma size, mean process length, or the number of process branching points (*H, n* = 5 control, *n* = 8 STZ). *I*: RNAseq data from retinal microglia taken from control and STZ-treated rats were screened for genes involved in the positive (GO: 0050729) and negative (GO:0050728) regulation of inflammation. While some inflammatory genes were altered, key inflammatory genes were unchanged after 4 weeks of diabetes. Data represented as mean ± SEM. * *p* < 0.05. Scale bar 50μm.

Previous work has shown blood-retinal barrier (BRB) integrity is compromised early in diabetes (38). Using vessel-dependent blood flow analysis (Fig.3*A*-*E*), we used the return to baseline after fluorescein peak (fluorescein offset) as a measure of BRB integrity. While no alteration in offset was observed for larger vessels, retinal capillaries showed a significant increase, indicative of fluorescein leakage / reduced BRB integrity (Fig. 4*F*; median regression analysis, *p* < 0.05). A breakdown in BRB can lead to immune cell infiltration and microglia activation, with microglial migration and morphological change indicative of classical activation observed in the retina, 1 month post-STZ (39). To assess whether altered microglial-vessel association occurred in the context of monocyte involvement / microglial activation, wholemounts were co-labelled with IB4 and Iba-1 and the number and morphology of microglia quantified in central and peripheral retina. Despite the increase in capillary fluorescein offset, there was no difference in the number of monocytes / retinal microglia (Fig. 4*G*; 2-way ANOVA, central *p* = 0.4, peripheral *p* = 0.9), or microglial morphology after 4 weeks of hyperglycaemia (Fig. 4*H*; 2-way ANOVA, cell body area *p* > 0.99, process length/cell *p* = 0.15, branch points/cell *p* > 0.99). Despite this, *Cx3cr1* expression was increased in the diabetic retina (*SI appendix*, Fig. S9). RNAseq analysis of microglial isolates from 4-week control and STZ-treated animals showed that of the 254 differentially expressed genes, 22 inflammatory response genes were identified, 15 of which were positive regulators (GO: 0050729), while 12 were negative regulators of inflammation (GO: 0050728) (Fig. 4*I*; *SI appendix,* tables S6 and S7). Importantly, chemokine and cytokines normally associated with microglial activation, including Tlr2, Il-1β, Cxcl10, TNF-a, IL-1a, C1q were not altered and there was no expression of the infiltrating monocyte marker gene, *Ccr2,* in our RNAseq dataset (40, 41). Thus, at this early stage of diabetes (4-weeks) when retinal capillaries are constricted, there is increased microglial-capillary interaction, which is independent of monocyte recruitment, classical microglial activation and a more generalised macroglial response.

### Microglial expression of vasoactive genes and control of capillary constriction are altered in early diabetes

To determine whether there was a loss of retinal vasomotor control during early diabetes, breathable oxygen was used to induce hyperoxic challenge and capillary diameter within the superficial vascular plexus was quantified using OCTA (Fig. 5*A* image). While control animals showed a distinct vasoconstriction in response to 100% oxygen, no constriction was observed in STZ-treated animals (Fig. 5*A*; 2-way ANOVA, *p* < 0.05). To explore whether this dysfunction was also evident in microglial-mediated vessel constriction, *ex vivo* retinal explants from control and STZ-treated animals (4 weeks post-STZ) were exposed to fractalkine and capillary diameter quantified. While constriction was evident in the control cohort, this response was absent in the STZ-treated animals (Fig. 5*B*, 2-way ANOVA, control *p* < 0.05, STZ-treated *p* = 0.99). When microglia were isolated from 4-week STZ-treated and control retinae and RNAseq performed, angiotensinogen (*Agt*) expression was increased 2.4 fold, while expression of the aryl hydrocarbon receptor gene (*Ahr*), a negative regulator of the RAS (42) was also increased (3.6 fold, Fig. 5*C*).

**Figure 5.**
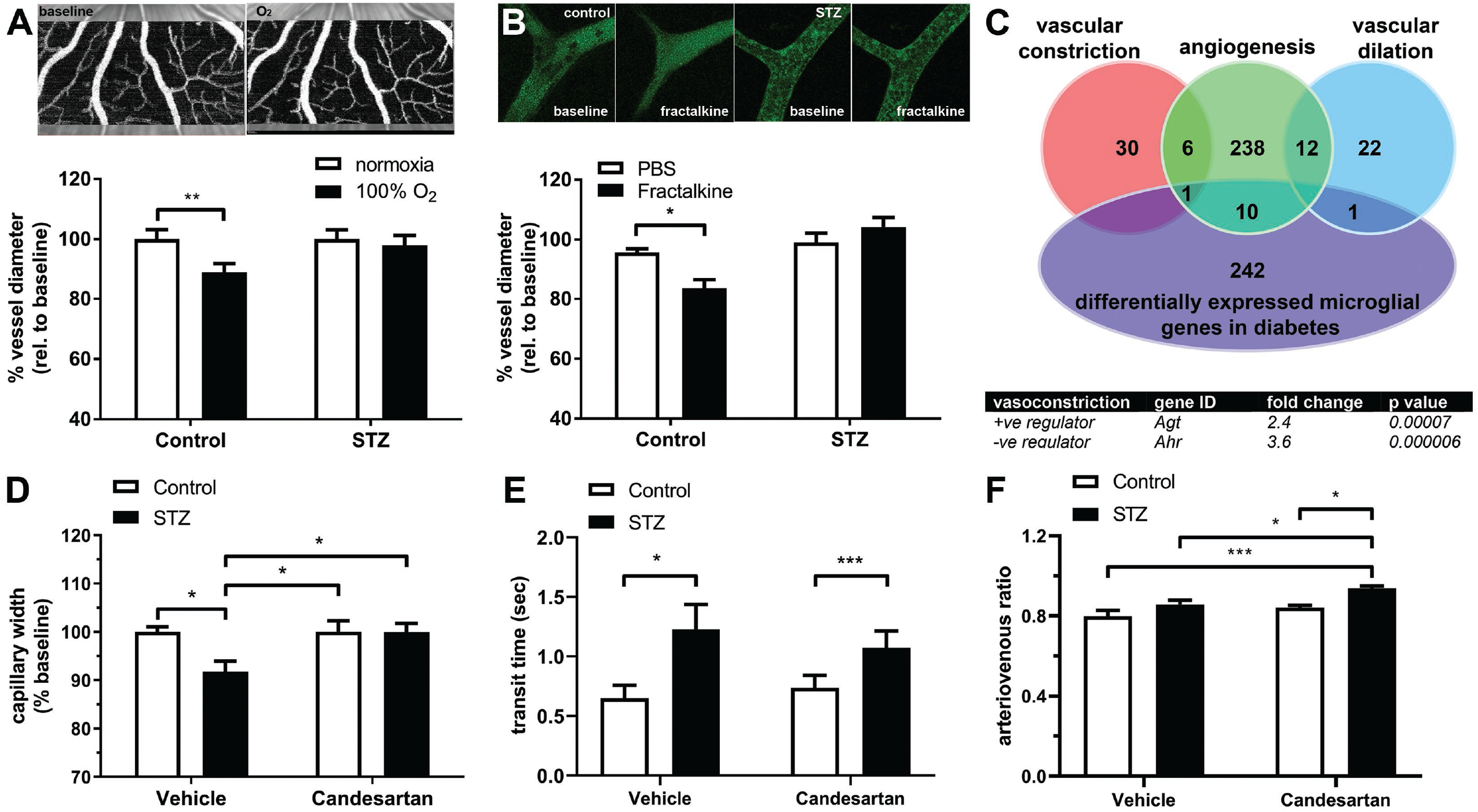
Vasoactive gene expression from retinal microglia and fractalkine-induced vasoconstriction are altered after 4 weeks of STZ-induced diabetes. *A*: The responsiveness of retinal vessels to hyperoxic challenge was explored *in vivo* using OCTA (*insets* show OCTA images from baseline and after exposure to O_2_). While hyperoxic challenge (filled bars) lead to constriction in the control group (*n* = 10 normoxia, *n* = 6 100% O_2_), no capillary constriction was observed in the STZ cohort (*n* = 12 normoxia, *n* = 7 100% O_2_). *B*: Microglial vasoregulation was investigated during diabetes, with 4-week STZ-treated and control retinae exposed to fractalkine *ex vivo* (representative control and STZ images in *inset*). While vessels from control retinae showed fractalkine-induced vasoconstriction (filled bar), STZ retinae exhibited no change in vessel diameter (*n* = 5 animals). *C:* Differential microglial gene expression data from 4 week control and STZ-treated animals were compared to vasomodulatory gene lists (vasoconstriction, GO:0097746; angiogenesis, GO:0001525; vasodilation, GO:0097746), with the RAS positive regulator angiotensinogen, (*Agt*) and negative regulator (*Ahr*) significantly dysregulated (FDR adjusted, citrate control *n* = 5, STZ n=4). *D*: OCTA was used to quantify retinal superficial capillary diameter in 4-week control and STZ-treated animals (unfilled and filled bars, respectively) exposed to candesartan or vehicle in their drinking water. In STZ-treated animals, capillary diameter returned to baseline in the candesartan-treated group (*n* = 7 control, *n* = 8, 5 STZ vehicle and candesartan, respectively). *E*: Retinal blood flow was quantified using arterio-venous transit time and showed increased transit time (slower blood flow) in STZ-treated animals independent of candesartan treatment (*n* = 8 control, *n* = 11 and 8 STZ vehicle and candesartan, respectively). *F*: Quantification of the arteriovenous ratio showed candesartan treatment increased the diameter of larger vessels in STZ-treated retinae relative to control and vehicle-treated tissues (*n* = 8 control, *n* = 11 and 8 STZ vehicle and candesartan, respectively). Data expressed as mean ± SEM, * *p* < 0.05, ** *p* < 0.01, \*\*\**p* < 0.001.

Based on the loss of vasomotor control in the diabetic retina and the dysregulation of the microglial RAS pathway, animals were rendered diabetic and treated with candesartan cilexetil or vehicle in their drinking water. At 4 weeks post-STZ, capillary diameter and retinal blood flow were quantified. OCTA analysis of superficial retinal capillaries showed a decrease in diameter within the vehicle control group, similar to that observed in Fig. 3I (Fig. 5*D*, 91.8 ± 2%, 2-way ANOVA, *p* < 0.05). This capillary constriction was not evident in STZ-treated animals exposed to candesartan, with diameters returning to control levels (Fig. 5*D*, 99.9 ± 1.8%, 2-way ANOVA, *p* > 0.99). However, despite this, retinal blood flow remained slower, with arterio-venous transit time increased in the vehicle and candesartan STZ-treated animals (Fig. 5*E*; median regression analysis *p* < 0.05, *p* < 0.001 respectively). Quantification of larger retinal vessels (arterioles and venules) showed systemic delivery of candesartan resulted in an increase arteriovenous ratio in the STZ-treated animals compared to candesartan-treated control (Fig. 5*F*; STZ 0.94 ± 0.01, control 0.84 ± 0.01, 2-way ANOVA p < 0.05) and vehicle-treated control and STZ animals (Fig. 5*F*; control 0.798 ± 0.03, STZ 0.86 ± 0.02, 2-way ANOVA p < 0.001 and 0.05, respectively).

Overall, these data show that in early diabetes, retinal vasomodulation is aberrant, with no evidence of microglial mediated vasoconstriction and specific dysregulation of the RAS. However, treatment with the AT1R inhibitor, candesartan, did not restore retinal blood flow, despite dilating the retinal capillaries.

## Discussion

The current study examined the role of microglia in local control of inner retinal blood supply. Microglia preferentially contact retinal capillaries that reside in the superficial vascular plexus, as well as contacting neuronal synapses within the inner retina. A novel role for microglia in vasomodulation within the retina and brain was identified, where addition of fractalkine induced capillary constriction. Subsequent characterisation within the retina showed this vasomodulation to be dependent on microglial contact and Cx3cr1 signalling. The microglial transcriptome contained gene transcripts for known vasoactive agents, while the AT1R inhibitor, candesartan, blocked capillary constriction, suggesting microglial vasoregulation likely occurs via modulation of local RAS. This was supported data showing fractakine-Cx3cr1-mediated upregulation of angiotensinogen. The microglial vasoregulatory role was further explored in the context of vascular dysfunction during early diabetes. After 4 weeks of experimental diabetes, retinal blood flow was reduced, coincident with constriction of the retinal capillaries within the superficial plexus and increased microglial-capillary association. However, there was no indication of classical microglial activation, nor a more generalised macroglial response during this early stage of diabetes. RNAseq data showed altered microglial expression of components of the RAS and there was a loss of microglial-mediated capillary constriction during diabetes. Finally, treatment with candesartan restored retinal capillary diameter in STZ-treated animals, however, retinal blood flow remained reduced.

### Microglial vasomodulation within the retina

The current data show that microglia are intimately associated with retinal vasculature, directly opposing pericytes and capillary areas free from pericytes, yet showing no particular preference for direct contact. Highlighting the functional significance of this interaction, stimulation of the microglial specific receptor Cx3cr1 via its sole ligand fractalkine, induced vasoconstriction, not only within the mouse and rat retina, but also in the brain. While the role of fractalkine-induced vessel constriction in the brain requires significantly more work to confirm microglial / Cx3cr1 involvement in areas exhibiting constriction, within the retina this effect was spatially discrete, occurring only in areas associated with microglial processes and was dependent on Cx3cr1 signalling, with *Cx3cr1^GFP/GFP^* retinae exhibiting no constriction, altered microglia-vessel contact and capillary diameter. These data directly implicate microglia in the capillary response to fractalkine. While previous work has identified microglia as a component of the blood-brain barrier (43), and involved in retinal and brain vascular development (44, 45), this is the first report of microglial-mediated vasomodulation. Furthermore, our data and those of others show microglia also monitor and modulate neuronal synapses during development, throughout adulthood and in response to activity (17, 46, 47), raising the possibility that microglia may contribute to neurovascular coupling, the process through which local blood flow is regulated by neuronal activity. As previous work in the retina suggests the existence of Müller cell-independent vasoregulatory mechanisms (11, 12), microglial vasoregulation may constitute one such alternative pathway, particularly within the superficial plexus. Further work exploring the structure of microglial-neuronal contact, it’s temporal characteristics and its response to altered neuronal activity will be required to properly characterise the role of microglia in the neurovascular unit.

### Microglial RAS involvement in capillary constriction

In order for microglia to directly mediate vessel constriction, they must express vasoactive factors. The RNAseq data from isolated retinal microglia highlighted several genes for vasoactive agents, including endothelin (*Edn1, 3)*, angiotensinogen (*Agt*) and arachidonate 5-lipoxygenase (*Alox5*), all of which are known to regulate retinal capillary tone (48). While retinal neuronal / glial cell contamination may confound the genes identified within the microglial isolate, the low levels of neuronal signature genes (Fig. S5) suggest any effect would be minor. Importantly, pre-incubation with the AT1R antagonist, candesartan, inhibited microglial-mediated vasoconstriction and incubation with fractalkine induced up regulation of microglial *Agt* expression which was not observed when Cx3cr1 was genetically ablated (*Cx3cr1^GFP/GFP^*). These data together with the dysregulated microglial genes identified during diabetes (*Agt* and *Ahr*), implicate the RAS in microglial-mediated vasoregulation. All components of the RAS have been observed within the retina, with angiotensin II (AngII) implicated in the vasoconstriction of all retinal vessels (arterioles, capillaries and venules) via AT1R (30, 31). While this microglial-mediated vasoregulation via the RAS is novel, microglia are known to express components of this pathway, including angiotensin converting enzyme, AT1R, AT2R (49). In addition to vessel constriction via the microglial RAS, microglial activation and inflammatory cytokine production has been described after AngII exposure within the brain and retina (50, 51). Thus, the modulation of the microglial RAS in normal tissue may be required for normal vessel control, whilst during pathology there may be a positive feedback cycle involving AngII, promoting microglial activation and inflammation.

Given the ultrastructural and immunocytochemical data suggesting microglia contact pericyte somata and processes, it is possible that microglia communicate directly with pericytes and utilise their vasomodulatory capacity (5) in order to constrict inner retinal capillaries. Supporting communication between both cell types, pericytes are able to modulate microglial phenotype during inflammation (52), while AT1R are expressed by pericytes enabling AngII-mediated constriction (31). In addition to pericytes, our data also show that microglia could elicit a response by communicating directly with endothelial cells (capillary areas free of pericytes), which are also known to express vasoregulatory substances (53). Finally, microglia may indirectly communicate with vessels via other retinal glia such as Müller cells, which express components of the RAS (54) and have been previously shown to regulate the inner retinal vasculature (9, 10). While a proposed mechanism is shown in Fig. 6, more work is required to explain how microglia signal to other members of the neurovascular unit to induce capillary constriction.

**Figure 6.**
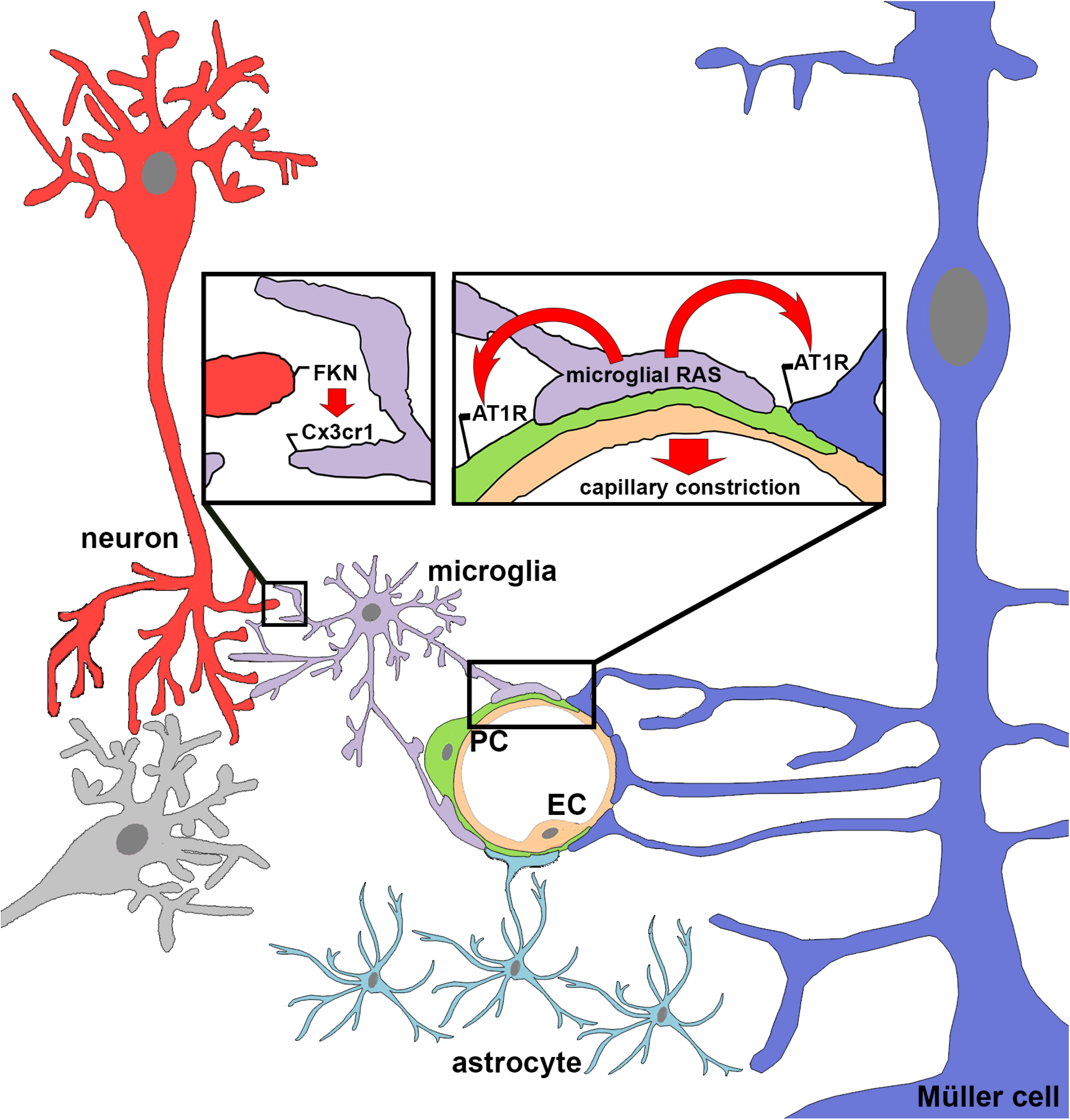
Schematic representation of microglial regulation of retinal capillary constriction. Data from this study shows microglia are structurally and functionally capable of involvement in the neurovascular unit. Microglia contact neuronal synapses and retinal capillaries (including pericytes) and activation of fractalkine-Cx3cr1 signalling results in capillary constriction, which is via an AT1R-dependent mechanism. Ultimately, capillary regulation may occur via direct microglial mechanism or may involve contributions from pericytes and / or Müller cells. FKN, fractalkine; RAS, renin angiotensin system; AT1R, angiotensin II receptor type 1; PC pericyte; EC, endothelial cell.

### Microglial involvement in capillary constriction during early diabetes and its effect on retinal blood flow

Our finding of reduced retinal blood flow throughout all retinal vessel types in response to short duration hyperglycaemia is supported by studies in both humans with diabetes and animal models of the disease (24). In contrast to larger retinal vessels which showed no alteration, a significant reduction in capillary diameter (~ −9%) within the superficial plexus was observed. To our knowledge this is a novel finding and while the change in capillary diameter is small, it would lead to large effect on blood flow, since capillaries constitute the majority of retinal vasculature (55). One estimate indicated a 6% dilation in capillary diameter (~0.32 μm) generated the majority of blood flow increase evoked by neuronal activity (5). In addition to static vessel change, retinal capillaries from STZ-treated animals failed to constrict after hyperoxic challenge. This is the first report of *in vivo* retinal capillary diameter measurement during vascular challenge, however, previous human studies have reported altered hyperoxic retinal vessel responses (blood flow) in patients with type 1 (56) and type 2 (57) diabetes.

As changes in the capillary network have been suggested to underlie the pathophysiology of early and later stage DR (24, 58, 59), it is tempting to speculate that microglial control of these vessels contribute to the vascular dysfunction in early diabetes. The data showing an increase in the number of microglial processes associated with the capillary network, the increase in microglial angiotensinogen (*Agt*) expression and the restoration of capillary diameter after candesartan cilexetil treatment all support this hypothesis. Even the increased microglial expression of aryl hydrocarbon receptor (*Ahr*), a negative regulator of vasoconstriction (42), may be incorporated into this theory, since recent work shows it contributes to vessel stiffness (60). Therefore, the increased *Ahr* and *Agt* expression may contribute to the phenotype of smaller and less responsive retinal blood vessels in early diabetes. Additional support for a microglial-specific effect on the retinal vasculature during diabetes comes from work undertaken in STZ-treated *Cx3cr1^GFP/GFP^* animals, which showed increased acellular capillaries after 4 months of hyperglycaemia (61). Further work using the STZ-treated *Cx3cr1^GFP/GFP^* model is required to specifically explore the capillary constriction evidenced early in diabetes.

The microglial dysregulation of the RAS suggests this pathway is altered in diabetes. These data are supported by our supplementary data (*SI appendix*, Fig. S9) and previous studies showing increased angiotensinogen within the vitreous of individuals with proliferative DR (62) and increased vitreal AngII concentrations and elevated retinal AngII, AT1R and AT2R levels in rodent models of diabetes (63, 64). As well as causing vasoconstriction, AngII is also known to uncouple pericytes from the endothelium, thereby altering vessel permeability and contributing to the development of microaneurysms, a key clinical determinant of DR (31). Validating the positive effects of candesartan on capillary vessel diameter and providing further support for the role of the RAS in DR, an earlier clinical trial showed candesartan blockade to be successful in preventing the onset of clinical grade DR in individuals with diabetes without DR (65). As these beneficial effects did not extend to preventing progression of DR in those with the disease, it suggests dysregulation of the RAS is relevant to the early, preclinical stage of DR.

Therefore, when the current data is considered together with the literature showing the RAS dysregulation during diabetes and the several studies showing increased fractalkine protein levels in the retina of STZ-treated rats (66, 67), a hypothesis can be formulated whereby in early diabetes, increased fractalkine expression together with enhanced microglial process-capillary interaction and a dysregulated microglial RAS, result in increased capillary vasoconstriction. While this potential role of microglial vasoregulation in DR is novel and unlike its inflammatory roles later in disease (39), further work is required to fully understand this early dysfunction and how it contributes to later pathology such as the diminished hyperaemic response observed in patients with diabetes (68, 69) and retinal hypoxia leading to later stage DR (36, 70).

While candesartan blockade did restore retinal capillary diameter to control levels in the current study, retinal blood flow remained decreased. This was surprising, as reversing capillary constriction would be expected to increase retinal blood flow, given the importance of the microvasculature (5, 55) and previous work showed candesartan cilexetil to restore blood flow in diabetic rats, all be it after 2 weeks post-STZ (71). However, quantification of arteriovenous ratio in the candesartan-treated STZ animals showed increased diameter of these larger vessels. These data, in conjunction with previous work which showed angiotensin II-dependent constriction of arterioles and venules (72), suggest that the dilation of the larger retinal vessels in the candesartan-treated STZ animals may result in reduced retinal blood velocity which masked the effect of the dilated capillaries. A more targeted delivery of factors for microglial RAS blockade may overcome these confounds and provide a clearer picture with respect to capillary dilation and retinal blood flow.

In summary, this study identifies a novel role for microglia in the modulation of capillaries within the CNS, particularly the retina. It highlights the involvement of the fractalkine-Cx3cr1 signalling axis and implicates the RAS in microglial-mediated capillary vasoregulation in the normal tissue and during the early stages of DR. While inhibition of the RAS pathway alters capillary constriction, it does not alter overall retinal blood flow in early diabetes. Further work investigating the cellular mechanism of microglial-induced vasoconstriction and intercellular signalling between microglia and other components of the neurovascular unit, will provide valuable information on the retinal vascular response in health and disease.

## Materials and Methods

### Animals

Animal procedures were approved by the University of Melbourne Ethics Committee (#1613867) and adhered to the National Health and Medical Research Council of Australia guidelines and the Guide for the Care and Use of Laboratory Animals. To explore the role of microglia in retinal vasomodulation, *Cx3cr1^GFP/+^* and *Cx3cr1^GFP/GFP^* mice were used which have one or both alleles of the monocyte-specific receptor, *Cx3cr1*, replaced with enhanced green fluorescent protein (EGFP) (73). To show that Cx3cr1 labels microglia within healthy retina and not infiltrating monocytes, immunohistochemistry was performed with select markers (*SI appendix*, Fig. S1). NG2-DsRed pericyte reporter mice were used to explore pericyte-microglial contact and were provided by Dr Sagdullaev. Adult mice were anaesthetised (ketamine:xylazine 67:13 mg/kg) and processed for transmission electron microscopy, live cell imaging or immunohistochemistry. Hyperglycaemia was induced in male adult (6 – 8-week-old) dark agouti rats via a single intraperitoneal injection of streptozotocin (STZ, 55 mg/kg, in trisodium citrate buffer, pH 4.5, Sigma-Aldrich Co, MO, USA), with control animals receiving an equivalent volume of vehicle. Blood glucose was measured 24 hours after injection to confirm conversion (>12 mmol/L; Accu-Chek Go, Roche Diagnostics, North Ryde, Australia). Weight and blood glucose levels were measured biweekly and STZ-treated animals received 2 units of insulin subcutaneously when blood glucose was ≥ 30 mmol/L (Novartis Pharmaceuticals Australia Pty. Ltd., North Ryde, Australia). A separate cohort of animals was treated with candesartan cilexetil (10 μg/ml; Sigma-Aldrich, #SML0245) or vehicle (PEG400 / Ethanol / Kolliphor® EL / water, 10:5:2:83, Sigma-Aldrich) in their drinking water, 24 hours after diabetes induction. After four weeks of diabetes, general anaesthesia was induced with an intraperitoneal injection of ketamine and xylazine (60 and 5 mg/kg respectively, Troy Laboratories Pty Ltd, Smithfield, Australia) prior to surgery, *in vivo* imaging and tissue isolation.

### Live cell imaging

Anesthetised *Cx3cr1^GFP/+^* and *Cx3cr1^GFP/GFP^* animals (n = 5, 6 respectively) were injected intraperitoneally with rhodamine B (Sigma-Aldrich) to label blood vessels, since IB4 labelling on live cell explants showed microglia cross reactivity (*SI appendix*, Fig. S4). After 5 minutes, animals were overdosed (pentabarbitone phosphate, 120 mg/kg) and retinae dissected into chilled Ames medium (Sigma-Aldrich) pre-bubbled with carbogen gas (95% O_2_, 5% CO_2_). Retinae were imaged on an inverted confocal microscope (Leica SP5), perfused with 37°C carbogenated Ames at 1ml/minute. Recombinant rat fractalkine (200 ng/ml; R&D Systems, MN, USA, #537-FT-025/CF) or vehicle (PBS) was introduced after 10 minutes of baseline recording and imaged for a further 10 minutes. At the end of this incubation, vessel diameter was measured at sites with or without microglial contact and measurements expressed as a percentage of baseline diameter of the same vessel region (taken as the average vessel diameter over the initial 10 minute baseline). *Ex vivo* preparations were imaged for a total of 30 minutes to limit vessel calibre variability. While this *ex vivo* preparation may have limitations with respect to retinal blood flow, all explants were treated identically and all effects were relative to initial baseline. The vascular response to fractalkine after 4 weeks of STZ-induced diabetes was measured using the above protocol, while to assess the role of the RAS in fractalkine induced constriction, *ex vivo* retinae were pre-incubated in Ames or Ames + 230 nM candesartan cilexetil (Sigma-Aldrich) for 10 minutes. Fractalkine (200ng/ml) was subsequently added and imaged for 10 minutes (n = 5 fractalkine + candesartan; n=7 fractalkine), at which time vessel diameter was quantified relative to pre-incubation baseline. While candesartan cilexetil is a prodrug that is generally activated during gastrointestinal absorption, carboxyl esterases are present within the retina (74) and our previous work shows candesartan cilexetil blocks angiotensin-induced vessel effects when delivered directly to the eye (51).

### *In vivo* video fluorescein angiography

For blood flow kinetic analysis of diabetic animals, *in vivo* video fluorescein angiography (VFA) was performed (n = 21 / group) as described previously using the Micron III rodent imaging system (Phoenix Research Labs, CA, USA) (33). This technique provides reliable quantification of blood flow kinetics using sodium fluorescein (1%, 100 μl/kg, Fluorescite 10%, Alcon Laboratories, NSW, Australia). Additional details are provided in the *SI appendix*. The time taken from fluorescein entry into the retina to half-maximum intensity (fill time), and the time taken to fall from maximum intensity to the midpoint between maximum and final intensity after 30 seconds of imaging (drain time) were recorded.

### *In vivo* Optical Coherence Tomography Angiography

To assess capillary diameter and capillary hyperoxic response *in vivo*, Optical Coherence Tomography Angiography (OCTA) was performed (OCT2 Spectralis, Heidelberg Engineering, Heidelberg, Germany). OCTA uses motion contrast imaging to generate real-time angiographic maps of the retinal vasculature (75). Volume scans (15 × 15-degree region of interest) were taken 2 – 3 disc diameters from the optic nerve. Each region consisted of 512 B-scans with each B-scan consisting of 512 A-scans. Superior and inferior retina were scanned in both eyes. The vascular response to hyperoxic conditions was measured in 4-week STZ-treated and control animals by exposing the animal to 100% oxygen via a nose cone (3 L/min). After a baseline image was taken, follow-up mode was used to acquire a second capillary image in the same retinal location, after 2 minutes of oxygen breathing.

### Immunocytochemistry

Rat or mouse retinae were processed for indirect immunofluorescence in wholemount or cross section, as previously described (76). Human tissue was obtained and processed as described previously (77). Retinal microglia were labelled with rabbit anti-ionized calcium-binding adapter molecule 1 (Iba-1,1:1000; Wako, Osaka, Japan) or expressed EGFP (*Cx3cr1^GFP/+^*, *Cx3cr1^GFP/GFP^*), while blood vessels were visualised with *Griffonia simplicifolia* isolectin B4 (IB4, FITC 1:75; Sigma-Aldrich; 647 fluorophore 1:100; Thermo Fisher Scientific, MA, USA). While IB4 has shown cross reactivity with brain microglia and activated retinal microglia (78, 79), we observe no cross reactivity in any fixed retinal tissues. We also show better vessel coverage using IB4 compared to the endothelial marker CD-31 (*SI appendix*, Fig. S10). Further details for immunolabelling are in *SI appendix*. All imaging was performed with a 20X objective on either Zeiss META / LSM800 confocals (Carl Zeiss, Oberkochen, Germany) or Leica SP5 (Wetzlar, Germany), while high resolution imaging of microglial-pericyte contact and EGFP expressing microglia was performed at 63X. For subsequent analysis of retinal wholemounts, tile scans were taken at the superficial vascular plexus with z-stacks (15.6μm) used to accommodate the variations in retinal mounting. All subsequent image analysis was performed on maximum intensity projections.

### Image analysis

#### Vessel morphology

Fundus images (n = 13 / group) were analysed for arteriole / venule width and tortuosity in MATLAB (Mathworks Inc., MA, USA) using the open source plugin ARIA (80) at an eccentricity of 1.5 and 2 disc diameters from the optic nerve. Capillary width (<15 μm) within the superficial vascular plexus (OCTA) was measured using AngioTool (81). Confocal wholemount images (n = 11 animals / group) were grouped into arterioles, venules and capillaries based on their corresponding VFA profile and vessel masks used to segment subsequent analysis in Metamorph (Molecular devices, CA, USA) using the angiogenesis tube formation application. Total vessel area was quantified in NIH ImageJ (82) for each vessel type and vessel density was expressed as percentage of vessel area covering the total retinal area. For all subjective measurements, individuals were blinded to the treatment group.

#### Microglial, glial, pericyte histology and vessel interaction

Microglia, pericytes and astrocytes from STZ-treated and control tissue were analysed in Metamorph utilising the neurite outgrowth application. Iba-1 positive microglia were segmented, counted and a mask generated. This microglial mask was overlaid on the vessel / pericyte masks and cells that overlapped with blood vessels by at least 0.82μm were considered touching and were calculated as a percentage of total cells. For microglial blood vessel and neuronal contact in *Cx3cr1^GFP/+^* and *Cx3cr1^GFP/GFP^* retinae, areas of colocalization between individual microglia and vessels and synapses were rendered as a 3D volume and expressed as a percentage of the total volume of the microglial cell (Imaris, Bitplane, Zurich, Switzerland; 3 microglia / quadrant / retina, n = 5 animals / genotype). To further characterise microglial-pericyte interaction a custom Metamorph script was used to quantify contacts (within 0.41μm) between microglia (Iba-1 positive) with pericyte somata and processes (NG2 positive), as well as capillary areas devoid of pericyte contact (NG2 negative, IB4 positive). Previous work has used EGFP in order to assess microglial contact with neurons in the retina and brain (17, 46, 83). Microglial morphology was also quantified using the automated neurite outgrowth application (Metamorph), while microglial-neuronal synapse and microglial-pericyte images were processed in Imaris. Astrocyte density within the ganglion cell layer was quantified for total retinal area and for overlap with each vessel type. Müller cell gliosis was quantified as previously described (84) (3 sections / animal, n = 6 animals).

### Electron microscopy

Pre-embedding immuno-electron microscopy was used to investigate the ultrastructural association of inner retinal capillaries and microglia in the *Cx3cr1^GFP/+^* retina, as previously described (76). The immunolabelling of EGFP using this protocol shows that EGFP is present close (<50nm) to the membrane at the tips of microglial processes (Fig. 1*C*).

### Microglial isolation and RNA-Seq

Retinae from control and 4-week STZ-treated rats (n = 5 control, n = 4 STZ, 12 weeks-old) were isolated, papain digested (Worthington Biochemical, NJ, USA), and labelled with CD11b-FITC conjugate (Miltenyi Biotec, Bergisch Gladbach, Germany) for microglial isolation (FACSAria III, BD Bioscience, San Jose, USA). RNA was isolated and RNAseq performed as in *SI appendix*. The identified microglial gene population was compared to other studies reporting microglial-enriched genes, as well as those detailing neuronal signature genes (*SI appendix,* Fig. S5). The RNAseq dataset was deposited into Gene Expression Omnibus (#GSE 139276). To explore fractalkine regulation of microglial RAS, retinae from C57bl6 and *Cx3cr1^GFP/GFP^* animals (n = 6) were incubated as above with fractalkine (200ng/ml, R&D Systems) or PBS for 2 hours at 37°C. Retinal microglia were isolated via FACS using the CD11b and EGFP labels. RNA was isolated and Smart-seq 2 performed with 13 cycles of pre-amplification followed by quantitative PCR (see *SI appendix*).

### Statistical analysis

Statistical significance was determined by two-tailed unpaired student’s t-test, two-way ANOVA or RM-ANOVA depending on the experiment (Prism 6.0, GraphPad, CA, USA). Where required a Tukey post-hoc analysis was performed. Blood flow analysis was undertaken using median regression analysis (STATA, StataCorp TX, USA). Alpha levels were set at 0.05. Numerical values are expressed as mean ± standard error of mean (SEM) unless otherwise stated.

## Acknowledgements

The authors thank Dr Leonid Churilov for his assistance with statistical analysis, Dr Christine Nguyen and Mr Darren Zhao for their assistance with blood gas analysis and Ms Satya Gunnam for her assistance with immunohistochemical staining and analysis. We also acknowledge the Biological Optical Microscopy and Cytometry Platforms at the University of Melbourne.

