## supplemental text and data for "Fractalkine-induced microglial vasoregulation occurs within the retina and is altered early in diabetic retinopathy"

**This PDF file includes:**

Supplementary text
Figures S1 to S10
Tables S1 to S7
Legends for Movies S1 to S2
SI References

**Other supplementary materials for this manuscript include the following:**

Movies S1 to S2

**Supplementary Information Text**

### Materials and Methods

#### Microglial process movement on retinal vessels

Dark agouti rats were anaesthetized, injected intraperitoneally with rhodamine B (Sigma-Aldrich) to label blood vessels and retinal explants established as described in the main text. Retinal microglia were labelled with Iba-1 and imaging performed on an inverted confocal microscope (Leica SP5). Baseline images were taken for 10 minutes, followed by the addition of PBS (10 minutes) and then either fractalkine or fractalkine + candesartan (10 minutes) using concentrations outlined in the main text. Select images were taken at the beginning, during and towards the end of each incubation (0, 4 and 8-minutes) and images were registered with reference to the landmark blood vessels. Retinal microglia from the 0, 4 and 8-minute images were pseudocoloured in NIH Image J  and images from 0 and 4 minutes (green and red microglia, respectively) and 4 -8 minutes (green and red microglia, respectively) overlayed to determine microglial process movement in early and later time periods during incubation.

#### Retinal blood flow velocity

After 4 weeks of STZ-induced diabetes, animals were anaesthetized, dilated and cannulated as described in the main text and placed on a temperature-controlled stage. Imaging of the retina was achieved after fluorescein injection using an Andor Neo 5.5 sCMOS camera (Scitech Pty Ltd, Preston Australia), enabling a high-speed video sequence (1700 frames per second) to be taken. *A*: A region of interest (500 x 100 pixels in size) was imaged in an area of retina containing artery and vein, superior to and 1-disc diameter away from the optic nerve. Each frame was registered to eliminate eye movement and the Canny Edge Detector used to identify vessel diameters (1). *B*: Blood vessel velocity was assessed on selected pixels by measuring their shift in pixel intensity over time at two defined distances within the vessel. Velocity could then be calculated using the highlighted equations. *C*: Grouped arteriole and venule velocities were calculated for control and STZ-treated animals showing a reduction in velocity in both vessel types in the diabetic animals ($n$ = 5). *D:* Arterio-venous transit time was calculated as the time between fluorescein appearing in the retinal arterioles and venules ($n$=25 control, $n$=22 STZ). Data presented as mean ± SEM and assessed using a 2-way ANOVA. *$p < 0.05$, ***$p < 0.001$.

### *In vivo* video fluorescein angiography and intraocular pressure, blood pressure and haematocrit calculation

After four weeks of diabetes, animals were anaesthetised (60 mg/kg ketamine and 5 mg/kg xylazine) and intraocular pressure was measured for STZ and control animals using a rebound Tonometer (Tonolab, iCare, Helsinki, Finland). An average from 10 readings per eye were taken for each animal ($n$ = 11). Animals received corneal anaesthetic (0.5% Alcaine, Alcon Laboratories), had their pupils dilated (Atropine 0.5%; Mydriacyl, Alcon Laboratories), while a femoral artery cannula was inserted and blood pressure was continuously monitored (LabChart, ADInstruments, Sydney, Australia) during Video Fluorescein Angiography ($n$ = 11). Arterial blood samples were also collected and total haemoglobin (ctHb) concentration quantified (ABL800 blood gas analyser, Radiometer, Copenhagen, Denmark), from which percentage haematocrit was calculated for control ($n$ = 17) and STZ-treated ($n$ = 18) cohorts (2). Data were expressed as mean ± SEM and assessed using a 2-way ANOVA. Translational image registration was applied to VFA videos to correct for eye movement, and masks were drawn to cover arterioles and venules. Capillary kinetics were characterised as the remaining retinal vasculature not covered by the arteriole or venule masks. The change in fluorescent intensity over time was calculated for every pixel in the image.

### Live cell imaging of brain vasculature

Preliminary experiments investigating the response of brain vasculature to fractalkine were performed in anesthetised dark agouti rats. The skull was exposed, thinned and imaging was performed on an upright wide field microscope using a 20X objective. Either fractalkine (200ng/ml, n = 3) or vehicle (PBS, n = 3) were administered by 10µl subdural injection and vessels imaged for 5 minutes. At multiple points along the blood vessel (<15µm), widths were measured at various times post-injection (NIH ImageJ; 5 second intervals out to 20 seconds, then 15 second intervals out to 320 seconds).

**Immunohistochemistry and validation of microglial Cx3cr1 expression in the retina**

In addition to the details supplied in the manuscript, human retinal sections mouse anti-vitronectin (1:100; Santa Cruz Biotechnology, TX, USA) was used to label blood vessels. Neuronal synapses were labelled with guinea pig anti-vesicular glutamate transporter 1 (VGLUT1, 1:500; Millipore, Bayswater, Australia) at the level of the inner plexiform layer, while pericytes were labelled with mouse anti-NG2 chondroitin sulphate proteoglycan (NG2, 1:1000; Millipore), Astrocytes were labelled with rabbit anti-glial fibrillary acidic protein (GFAP, 1:10,000; Dako, Santa Clara, CA, USA) and imaged only in the ganglion cell layer. Gliotic Müller cells were quantified by co-labelling retinal cross sections with rabbit anti-GFAP and mouse anti-glutamine synthetase (GS, 1:1000; Millipore). Cell nuclei were labelled with 4′,6-diamidino-2-phenylindole (DAPI). Secondary antibodies were all raised in goat to the specific primary host (Alexa Goat anti-rabbit 594/488, anti- mouse 594/488, Thermo Fisher Scientific). For retinal pericyte density imaging was performed with a 20X objective on either Zeiss META confocal (Carl Zeiss, Oberkochen, Germany) or Leica SP5 (Wetzlar, Germany). Pericyte density was quantified per vessel area in central and peripheral retina (*n* = 11). Data were expressed as mean ± SEM and analysed using a 2-way ANOVA.

To determine whether eGFP expression in the normal retina specifically labels microglia, fixed Cx3cr1$^{+/GFP}$ retinae were double labelled with select markers to distinguish infiltrating monocytes from resident microglia (see immunocytochemistry in materials and methods). Primary antibodies to anti-ionized calcium-binding adapter molecule 1 (Iba-1,1:1000; Wako, Osaka, Japan), the purinergic receptor, P2Y12 receptor (P2Y12R, 1:500; Alaspec, Fremont, CA, USA) were used. Flow cytometry (FACSAria III, BD Bioscience, San Jose,

USA) was also used and the extent of GFP- and C-C chemokine receptor type 2- (CCR2, BioLegend, San Diego, USA) and integrin αM-labelled cells (CD11b-FITC conjugate, Miltenyi Biotec, Bergisch Gladbach, Germany) quantified, n=4. In order to further assess microglial-pericyte contact, NG2-DsRed reporter mice were labelled with Iba1 (as in materials and methods) and CD31 (1:8000, R&D Systems). Images were taken on a Leica SP8 confocal microscope using 63x oil objective and were rendered with Imaris. For assessment of retinal blood vessels, retinal wholemounts were labelled with IB4 (as in materials and methods), EGFP (microglia) and CD31 (endothelial cells, as above).

**RNAseq and gene expression analysis**

For RNAseq of microglial isolates, total RNA was extracted from isolated populations (RNeasy Micro Kit, Qiagen, Hilden, Germany) and purity analysed (Agilent Technologies, Santa Clara, CA, USA). A SmartSeq v4 kit (Clontech Laboratories, Mountain View, CA, USA) was used for a preamplification step. The Australian Genome Research Facility performed 50 bp single end reads at a depth of 19 – 34 million reads/sample using Illumina HiSeq (San Diego, CA, USA), before mapping the identified constructs to the rat genome and calculating differential expression between control and STZ groups.

For quantitative PCR analysis of fractalkine incubated C57bl6 and *Cx3cr1*$^{GFP/GFP}$ retinae, rat specific primers (*Agt*: FW 5′-ttgggtgctgaggcaaatct-3′, antisense: 5′-ccacattttgggggttat-3′; *Hprt*: FW 5′-cctaaaacacagcggcaagttgaa-3′, antisense: 5′-ccacaggactagaacgtctgctag-3′; *Gapdh*: FW 5′-tgtatccgttgtggatctga-3′, antisense: 5′-ttgctgttgaagtcacaggag-3′) were used to produce cDNA products which were purified and converted into cRNA (Megascript, Thermo Fisher Scientific). A four-point standard curve for *Agt*, *Hprt*, *Gapdh* was included in pre-amplification and used to quantify with gene copies relative to housekeeping genes, *Hprt* and *Gapdh*. For gene expression in whole retina, total RNA was isolated from dissected control (vehicle and candesartan, *n* = 8 each) and STZ-treated (vehicle and candesartan, *n* = 8 each) rat retinae, reverse transcribed (Tetro, Bioline, London,

UK) and amplified (Sensifast SYBR, Bioline) using the Rotorgene 3000 (Qiagen, Hilden, Germany). Again, gene expression was quantified relative to gene specific standards and the housekeeping gene *Gapdh* and expressed as copy of gene of interest / copy housekeeper. (Cx3cr1 Fw ggccttgagcgacctgctctttg, Cx3cr1 Rv gatgctgatgacggtgatgaagaa; At1r Fw tcacctgcatcatcatctgg, At1r_Rv agctggtgagaatgataagg; Gapdh Fw tgtatccgttgtggatctga; Gapdh Rv ttgctgttgaagtcacaggag). All data were assessed using a 2-way ANOVA with a Bonferroni post hoc analysis (GraphPad Prism).

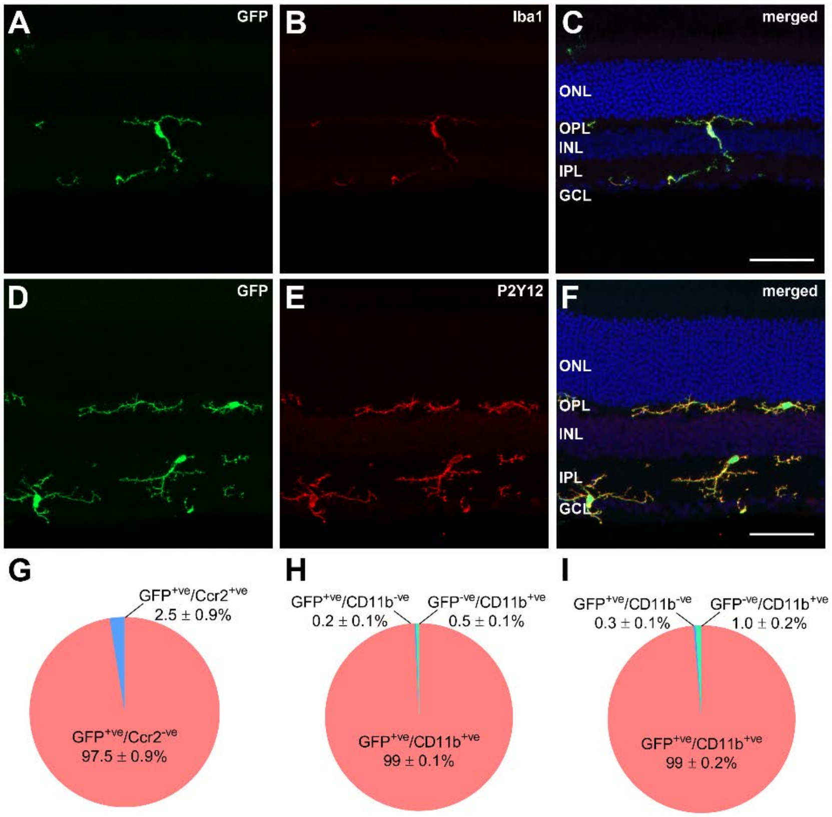


**Fig. S1. GFP expression in the *Cx3cr1$^{GFP/GFP}$* and *Cx3cr1$^{+/GFP}$* retinae are limited to resident microglia.** *Cx3cr1$^{GFP/GFP}$* and *Cx3cr1$^{+/GFP}$* retinae were labelled with select markers to determine the specificity of EGFP labelling. *A-C:* Double labelling with EGFP (*A*) and Iba-1 (*B*) within the retina shows 100% co-localisation (*C*), discounting the presence of infiltrating monocytes, which are Iba1-negative (3) and also supporting our use of this label in quantifying microglial-vascular interactions (Fig. 4). *D-F:* The purinergic receptor, P2Y12R, has been reported to be a "signature receptor" for microglia (4) and not found on macrophages (5). P2Y12 (*D*) and EGFP (*E*) show 100% co-localisation (*F*) suggesting these cells are in fact, microglia. *G:* This was also supported by via flow cytometry showing little co-localisation of CCR2 expression on the EGFP cells (GFP$^{hi}$CCR2$^{hi}$) which is found on infiltrating monocytes (6). *H-I*: We further show that >99% of the retinal EGFP labelled cells also label with CD11b (Cx3cr1$^{+/GFP}$ and Cx3cr1$^{GFP/GFP}$ respectively), which we used for our microglial isolation for our RNAseq study. Flow cytometry data shown as mean ± SEM, n=3 (*G*), n=4 (*H, I*). Scale 50µm

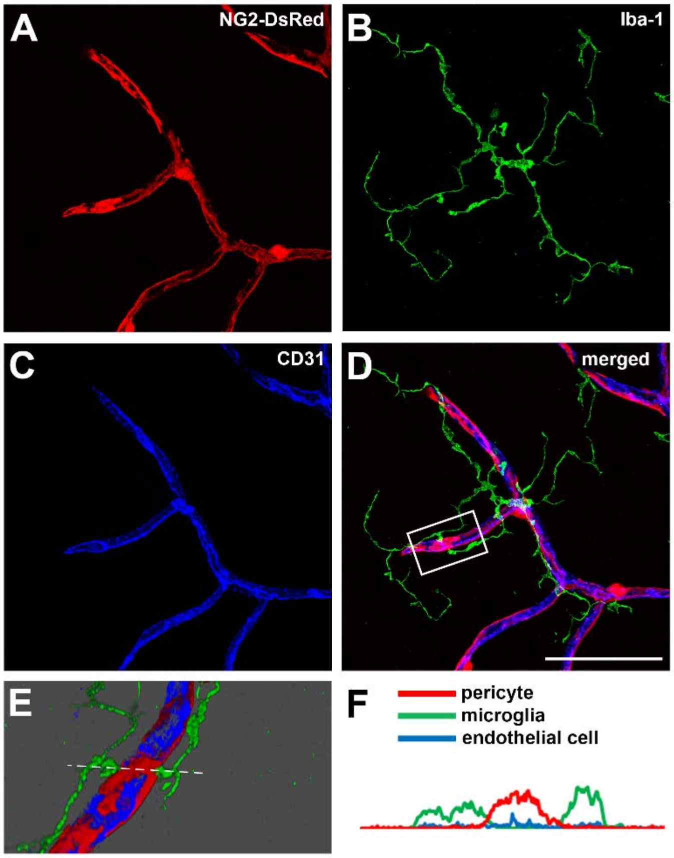


**Fig, S2. Microglia make contact with retinal pericytes on capillaries.** Retinal wholemounts from NG2-DsRedreporter mouse were used to assess microglial-pericyte contact. *A-D:* Pericytes were labeled by DsRed under the control of the NG2 promoter, red (*A*) and were stained for Iba-1 (*B*, microglia, green), and CD31 (*C*, endothelial cells, blue). *D:* In the merged image, a microglial process is observed to make contact with a pericyte (square in *D*). *E:* An Imaris rendering of the area showing microglial-pericyte contact. *F:* taking a cross section through the proposed contact area (dotted line in *E*), the traces show direct contact between the pericyte (red line) and microglial process (green line). Scale bar 50μm.

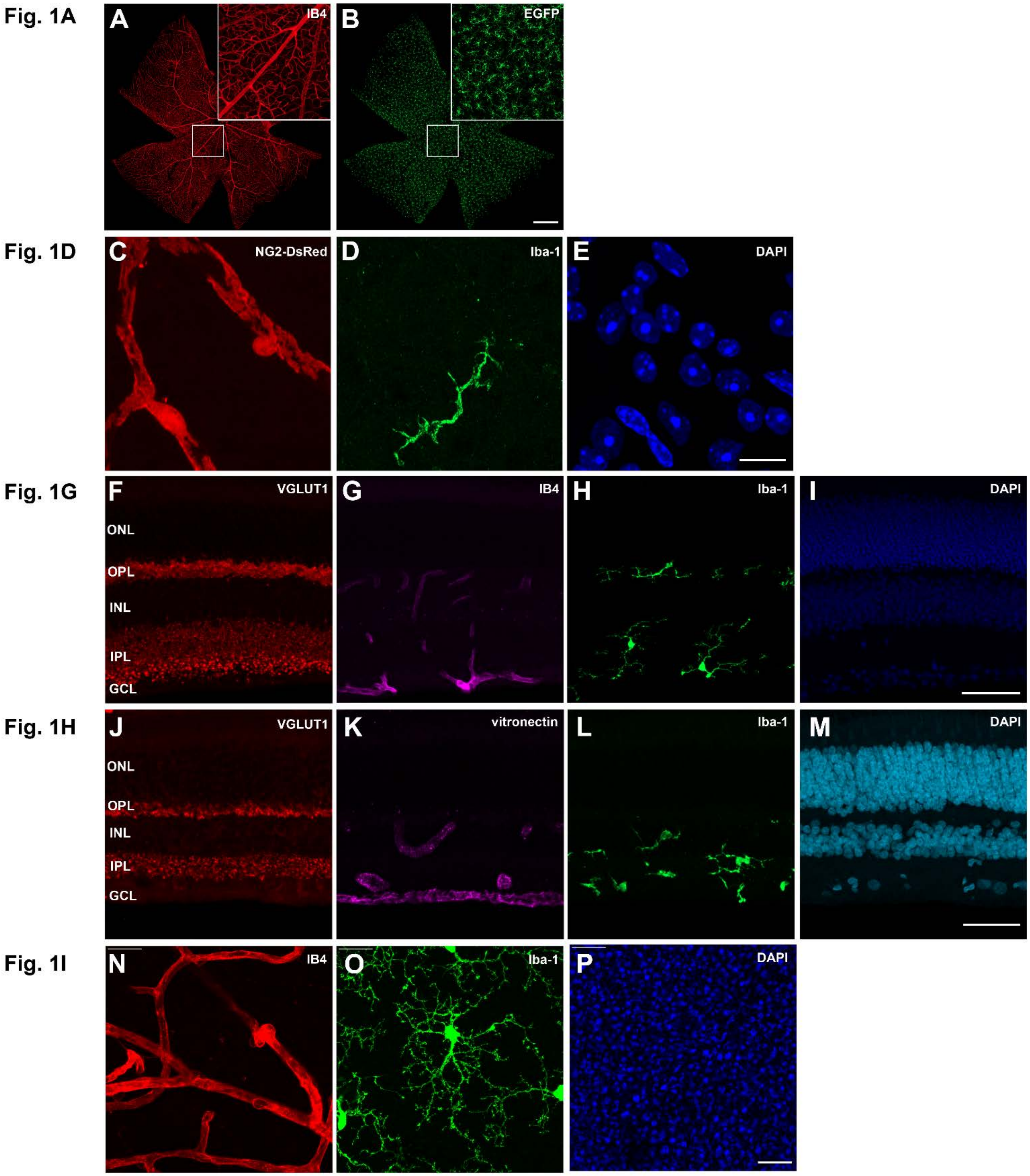


**Fig. S3. Individual imaging channels for immunohistochemical detection in Fig.1.** Each individual imaging channel and label for the merged images in Fig. 1 of the manuscript is shown. The scale bars are *A*,*B* 500 µm, *C*-*E* 10µm, *F*-*M* 50µm, *N*-*P* 20µm.

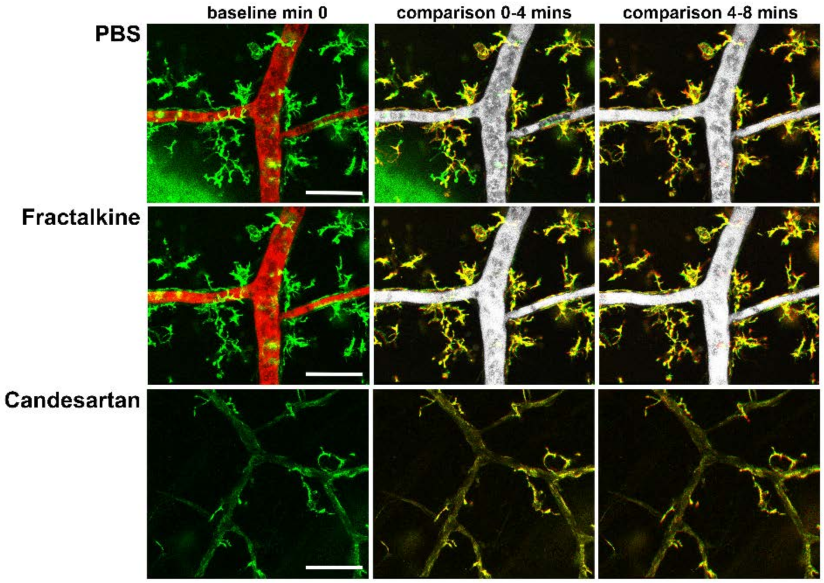


**Fig. S4. Retinal microglial processes show limited movement at the vascular surface.** *Ex vivo* retinal preparations (dark agouti rats) were isolated and imaged (Leica SP5; IB4, microglia green; rhodamine B, blood vessels red / grey scale). Explants were perfused with PBS for approximately 10 minutes, prior to the addition of recombinant rat fractalkine (200 ng/ml) or fractalkine + candesartan cilexetil (230nM). Representative images are shown at the start of the PBS, fraktaline and fractalkine + candesartan additions (baseline min 0), while subsequent images at 4 minutes were compared back the baseline, while the 8-minute image was compared to the 4-minute time point to gauge microglial process movement (red / green is evidence of process movement, yellow indicates a static process). As can be observed in the PBS and fractalkine comparisons, there is limited microglial process movement as evidenced by the colocalization (few red or green processes). Blood vessels were represented in grey scale in the comparison panels and the candesartan images were taken from a series where no rhodamine B was injected (IB4 labelled both microglia and retinal vessels).

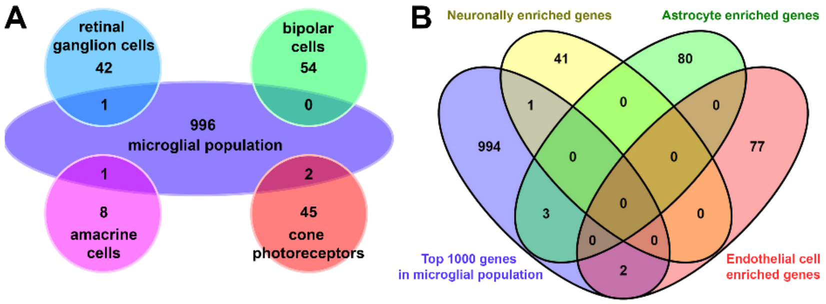


C

| Reference | Analysis type | Number of genes | Present in microglial population | Present in top 1000 expressed genes |
|---|---|---|---|---|
| Gyoneva et al | highest expression TPM* | 50 | 42 (84%) | 38 (76%) |
| Zheng et al | highest expression FPKM† | 50 | 46 (92%) | 40 (80%) |
| Butovsky et al | enriched compared to other brain cells | 86 | 77 (90%) | 46 (53%) |
| Hickman et al | microglial sensome – transmembrane proteins | 50 | 39 (78%) | 31 (62%) |

**Fig. S5. CD11b isolated microglial gene population shows limited neuronal gene contamination and reflects previously published microglial signature genes.** *A:* A comparison was performed between previously published data highlighting neuronally-enriched genes for retinal ganglion (7), amacrine (8), bipolar (9) and cone photoreceptors (10). The comparison showed limited expression of these neuronal genes within the CD11b isolated microglial population. B: A similar comparison was performed on neuronal, astrocyte and endothelial cell genes from published work in the brain (11), with again limited contamination observed. C: The genes from the CD11b isolated microglial population were compared to several studies reporting microglial-enriched genes and showed significant correlation (4,11,12,13).

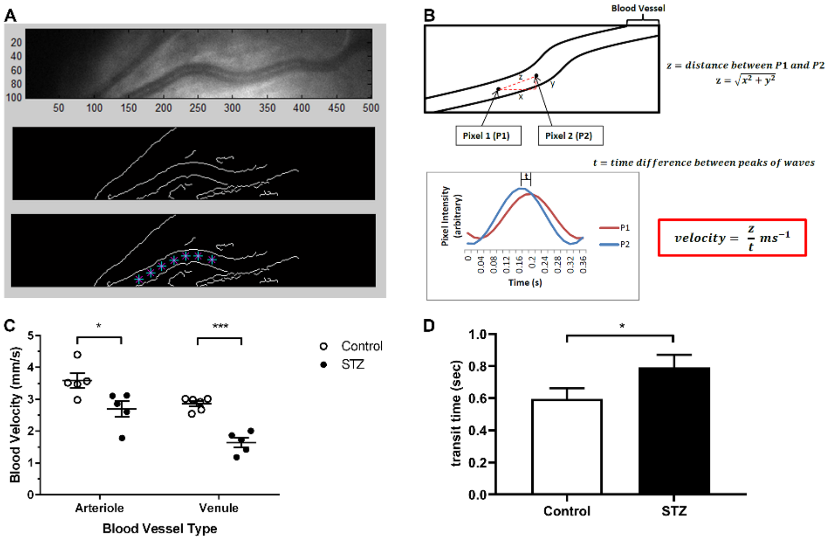


**Fig. S6. Retinal arteriole and venule blood flow velocity was slower and transit time increased in STZ-treated animals.** High speed retinal imaging was performed on control and STZ-treated animals to measure blood velocity. *A*: A representative still image from the high-speed video showing a retinal arteriole and the resultant border detection method used to isolate the vessel and find its luminal centre. *B*: The pixel intensity of two selected points along the vessel were analysed, and a sinusoidal curve fitted. The phase shift between these two fitted curves was used to detect the time taken for blood to travel the known distance (z), allowing velocity to be calculated. *C*: Compared to controls, STZ-treated animals showed a significant reduction in blood velocity in both venules and arterioles ($n$ = 5. *$p < 0.05$, ***$p < 0.001$). *D*: Arterio-venous transit time was calculated as the time between fluorescein appearing in the retinal arterioles and venules ($n$=25 control, $n$=22 STZ. *$p$<0.05). Data presented as mean ± SEM.

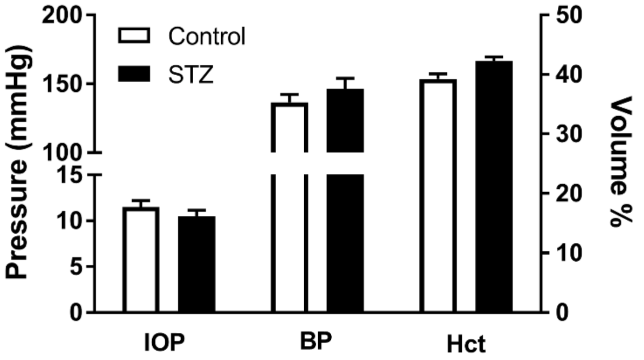


**Fig. S7. Intraocular pressure, blood pressure and haematocrit are not altered in STZ-treated animals.** Factors known to affect blood flow systemically and locally were unchanged after 4 weeks of STZ-induced diabetes. No difference was observed in intraocular pressure (IOP, *n* = 11), systolic blood pressure (BP, *n* = 11), or calculated haematocrit (Hct, *n* = 17 control, *n* = 18 STZ) between control (unfilled bars) and STZ-treated (filled bars) animals. Group data expressed as mean ± SEM.

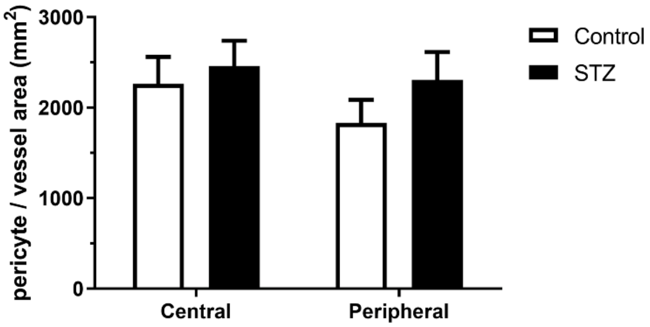


**Fig. S8. Pericyte density is not altered in STZ-treated animals.** Pericyte numbers were quantified in the central and peripheral retina and expressed relative to retinal vessel area. No difference was observed in pericyte density between control (*n* = 11, unfilled bars) and STZ-treated (*n* = 11, filled bars) animals. Group data expressed as mean ± SEM.

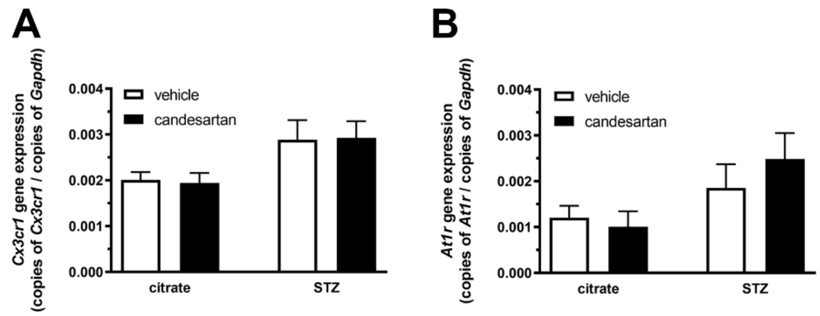


**Fig. S9. Retinal *Cx3cr1* and *At1r* gene expression are increased in STZ-treated animals.** Total retinal RNA was isolated and qPCR performed to quantify *Cx3cr1* and *At1r* gene expression relative to the housekeeping gene *Gapdh*. STZ-induced diabetes resulted in a significant increase in *Cx3cr1* and *At1r* gene expression (2-way ANOVA, $p<0.01$ and $p < 0.05$ respectively) in STZ-treated animals ($n = 8$) compared to control ($n = 8$). There was no effect of candesartan treatment (filled bars) compared to vehicle (unfilled bars). Group data expressed as mean ± SEM.

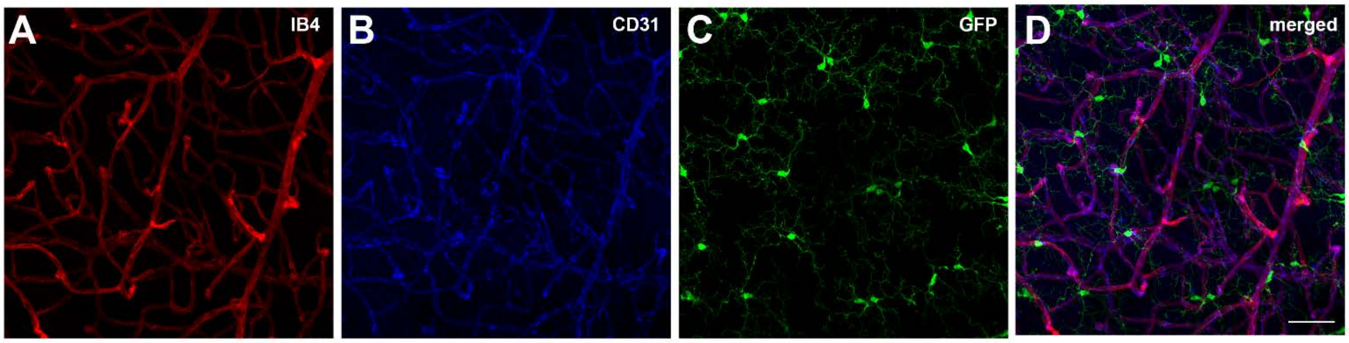


**Fig. S10. Blood vessel labelling with IB4 shows no microglial cross reactivity in the retina and shows improved vessel coverage compared to CD31.** $Cx3cr1^{GFP/+}$ mouse retina were fixed as described in the materials and methods and labelled with the blood vessel marker IB4, the endothelial cell marker CD31, with colocalization undertaken with EGFP (microglia). *A-D:* IB4 (*A*) and CD31 (*B*) show select labelling of the retinal vasculature, with no evidence of microglial cross reactivity (compare with EGFP in *C* and merged image in *D*). Scale 50µm

**Table S1: Microglial signature genes detected in FACS isolated retinal microglia.** Using flow cytometry and deep RNA sequencing a list of specific microglial marker genes was compiled by Chiu et al. (29). When the RNAseq results from our FACS retinal microglia were compared to this list, 23/29 microglial signature genes were identified in the retinal microglial population.

| **Gene ID** | **Gene name** | **Detected** |
|---|---|---|
| Adora3 | adenosine A3 receptor | No |
| Bco2 | beta-carotene oxygenase 2 | Yes |
| Capn3 | calpain 3 | Yes |
| Ccl3 | chemokine (C-C motif) ligand 3 | Yes |
| Ccl4 | chemokine (C-C motif) ligand 4 | Yes |
| Ccrl2 | Chemokine C-C motif chemokine receptor like 2 | Yes |
| Csmd3 | CUB and Sushi multiple domains 3 | No |
| Cx3cr1 | Chemokine CX3C motif receptor 1 | Yes |
| Egr1 | early growth response 1 | Yes |
| F11r | F11 receptor | Yes |
| Fosb | FosB proto-oncogene, AP-1 transcription factor subunit | Yes |
| G530011006Rik | unknown | No |
| Gal3st4 | galactose-3-O-sulfotransferase 4 | Yes |
| Gm10790 | predicted gene 10790 | No |
| Golm1 | golgi membrane protein 1 | Yes |
| Gpr84 | G protein-coupled receptor 84 | Yes |
| Grap | GRB2-related adaptor protein | Yes |
| Gtf2h2 | general transcription factor IIH subunit 2 | Yes |
| IL21r | interleukin 21 receptor | Yes |
| Lag3 | lymphocyte activating 3 | Yes |
| Lrrc3 | leucine rich repeat containing 3 | No |
| Olfml3 | olfactomedin-like 3 | Yes |
| P2ry13 | Purinergic receptor P2Y G protein-coupled 13 | Yes |
| Ptgs1 | prostaglandin-endoperoxide synthase 1 | Yes |
| Siglec H | sialic acid binding Ig-like lectin H | No |
| Slc2a5 | Solute carrier family 2 member 5 | Yes |
| Slco2b1 | solute carrier organic anion transporter family, member 2b1 | Yes |
| Tagap | T-cell activation RhoGTPase activating protein | Yes |
| Tmem119 | Transmembrane protein 119 | Yes |

**Table S2: Microglial genes involved in angiogenesis.**

Genes expressed by retinal microglia were compared against genes involved in angiogenesis (GO:0001525, filtered for Rattus norvegicus). Of the 407 rat genes probed, 268 microglial genes were identified as having a role in angiogenesis.

| Gene ID | Gene name | Vasoactive action |
|---|---|---|
| Ackr3 | atypical chemokine receptor 3 | angiogenesis |
| Acvr1 | activin A receptor type 1 | branching involved in blood vessel morphogenesis |
| Acvrl1 | activin A receptor like type 1 | positive regulation of angiogenesis |
| Adam15 | ADAM metallopeptidase domain 15 | angiogenesis |
| Adam8 | ADAM metallopeptidase domain 8 | angiogenesis |
| Add1 | adducin 1 | positive regulation of angiogenesis |
| Adgra2 | adhesion G protein-coupled receptor A2 | angiogenesis |
| Adgrb2 | adhesion G protein-coupled receptor B2 | negative regulation of angiogenesis |
| Adgrg1 | adhesion G protein-coupled receptor G1 | angiogenesis |
| Adipor2 | adiponectin receptor 2 | vascular wound healing |
| Adm | adrenomedullin | positive regulation of angiogenesis |
| Adrb2 | adrenoceptor beta 2 | negative regulation of angiogenesis |
| Aggf1 | angiogenic factor with G patch and FHA domains 1 | positive regulation of angiogenesis |
| Agt | angiotensinogen | negative regulation of angiogenesis |
| Ahr | aryl hydrocarbon receptor | branching involved in blood vessel morphogenesis |
| Aimp1 | aminoacyl tRNA synthetase complex-interacting multifunctional protein 1 | negative regulation of angiogenesis |
| Alox12 | arachidonate 12-lipoxygenase, 12S type | positive regulation of angiogenesis |
| Angpt2 | angiopoietin 2 | angiogenesis |
| Angpt4 | angiopoietin 4 | angiogenesis |
| Angptl4 | angiopoietin-like 4 | angiogenesis |
| Anpep | alanyl aminopeptidase, membrane | angiogenesis |
| Anxa2 | annexin A2 | angiogenesis |
| Anxa3 | annexin A3 | positive regulation of angiogenesis |
| Apold1 | apolipoprotein L domain containing 1 | angiogenesis |
| Arhgap24 | Rho GTPase activating protein 24 | angiogenesis |
| Atp5b | ATP synthase, H+ transporting, mitochondrial F1 complex, beta polypeptide | angiogenesis |
| B4galt1 | beta-1,4-galactosyltransferase 1 | angiogenesis involved in wound healing |
| Brca1 | BRCA1, DNA repair associated | positive regulation of angiogenesis |
| Btg1 | BTG anti-proliferation factor 1 | positive regulation of angiogenesis |
| C1galt1 | core 1 synthase, glycoprotein-N-acetylgalactosamine 3-beta-galactosyltransferase, 1 | angiogenesis |
| C3 | complement C3 | positive regulation of angiogenesis |
| C3ar1 | complement C3a receptor 1 | positive regulation of angiogenesis |

| | | |
|---|---|---|
| C5ar1 | complement C5a receptor 1 | positive regulation of angiogenesis |
| C6 | complement C6 | positive regulation of angiogenesis |
| Calcrl | calcitonin receptor like receptor | angiogenesis |
| Casp8 | caspase 8 | angiogenesis |
| Cav1 | caveolin 1 | angiogenesis |
| Ccl12 | chemokine (C-C motif) ligand 12 | angiogenesis |
| Ccl2 | C-C motif chemokine ligand 2 | negative regulation of angiogenesis |
| Ccl24 | C-C motif chemokine ligand 24 | positive regulation of angiogenesis |
| Ccr3 | C-C motif chemokine receptor 3 | positive regulation of angiogenesis |
| Ccr5 | chemokine (C-C motif) receptor 5 | positive regulation of cell migration involved in sprouting angiogenesis |
| Cd34 | CD34 molecule | positive regulation of angiogenesis |
| Cd36 | CD36 molecule | negative regulation of angiogenesis |
| Cdc42 | cell division cycle 42 | sprouting angiogenesis |
| Cdh13 | cadherin 13 | sprouting angiogenesis |
| Ceacam1 | carcinoembryonic antigen related cell adhesion molecule 1 | regulation of sprouting angiogenesis |
| Chi3l1 | chitinase 3 like 1 | positive regulation of angiogenesis |
| Cib1 | calcium and integrin binding 1 | positive regulation of cell migration involved in sprouting angiogenesis |
| Clic4 | chloride intracellular channel 4 | angiogenesis |
| Col18a1 | collagen type XVIII alpha 1 chain | angiogenesis |
| Col4a1 | collagen type IV alpha 1 chain | branching involved in blood vessel morphogenesis |
| Col4a2 | collagen type IV alpha 2 chain | negative regulation of angiogenesis |
| Cspg4 | chondroitin sulfate proteoglycan 4 | angiogenesis |
| Ctgf | connective tissue growth factor | angiogenesis |
| Ctnnb1 | catenin beta 1 | negative regulation of angiogenesis |
| Ctsh | cathepsin H | positive regulation of angiogenesis |
| Cx3cl1 | C-X3-C motif chemokine ligand 1 | positive regulation of angiogenesis |
| Cx3cr1 | C-X3-C motif chemokine receptor 1 | regulation of angiogenesis |
| Cxcl10 | C-X-C motif chemokine ligand 10 | negative regulation of angiogenesis |
| Cxcl12 | C-X-C motif chemokine ligand 12 | branching involved in blood vessel morphogenesis |
| Cxcr2 | C-X-C motif chemokine receptor 2 | positive regulation of angiogenesis |
| Cxcr3 | C-X-C motif chemokine receptor 3 | angiogenesis |
| Cxcr4 | C-X-C motif chemokine receptor 4 | branching involved in blood vessel morphogenesis |
| Cybb | cytochrome b-245 beta chain | positive regulation of angiogenesis |
| Cyr61 | cysteine-rich, angiogenic inducer, 61 | positive regulation of angiogenesis |
| Cysltr1 | cysteinyl leukotriene receptor 1 | positive regulation of angiogenesis |
| Dab2ip | DAB2 interacting protein | negative regulation of angiogenesis |
| Dag1 | dystroglycan 1 | angiogenesis involved in wound healing |
| Ddah1 | dimethylarginine dimethylaminohydrolase 1 | positive regulation of angiogenesis |

| | | |
|---|---|---|
| Dicer1 | dicer 1 ribonuclease III | angiogenesis |
| E2f8 | E2F transcription factor 8 | sprouting angiogenesis |
| Ecm1 | extracellular matrix protein 1 | positive regulation of angiogenesis |
| Edn1 | endothelin 1 | branching involved in blood vessel morphogenesis |
| Efna1 | ephrin A1 | angiogenesis |
| Efnb2 | ephrin B2 | cell migration involved in sprouting angiogenesis |
| Egfl7 | EGF-like-domain, multiple 7 | angiogenesis |
| Egr3 | early growth response 3 | cell migration involved in sprouting angiogenesis |
| Eif2ak3 | eukaryotic translation initiation factor 2 alpha kinase 3 | angiogenesis |
| Elk3 | ELK3, ETS transcription factor | angiogenesis |
| Eng | endoglin | positive regulation of angiogenesis |
| Enpep | glutamyl aminopeptidase | angiogenesis |
| Epas1 | endothelial PAS domain protein 1 | angiogenesis |
| Epha2 | Eph receptor A2 | negative regulation of angiogenesis |
| Ephb1 | Eph receptor B1 | angiogenesis |
| Ephb3 | Eph receptor B3 | angiogenesis |
| Erap1 | endoplasmic reticulum aminopeptidase 1 | positive regulation of angiogenesis |
| Ets1 | ETS proto-oncogene 1, transcription factor | positive regulation of angiogenesis |
| F3 | coagulation factor III, tissue factor | positive regulation of angiogenesis |
| Fbxw7 | F-box and WD repeat domain containing 7 | regulation of cell migration involved in sprouting angiogenesis |
| Fgf1 | fibroblast growth factor 1 | positive regulation of angiogenesis |
| Fgf2 | fibroblast growth factor 2 | positive regulation of angiogenesis |
| Fgf9 | fibroblast growth factor 9 | angiogenesis |
| Fgfr1 | Fibroblast growth factor receptor 1 | angiogenesis |
| Flna | filamin A | angiogenesis |
| Flt1 | FMS-related tyrosine kinase 1 | angiogenesis |
| Fn1 | fibronectin 1 | angiogenesis |
| Fzd5 | frizzled class receptor 5 | angiogenesis |
| Gna13 | G protein subunit alpha 13 | angiogenesis |
| Gpi | glucose-6-phosphate isomerase | angiogenesis |
| Gpld1 | glycosylphosphatidylinositol specific phospholipase D1 | cell migration involved in sprouting angiogenesis |
| Gpx1 | glutathione peroxidase 1 | angiogenesis involved in wound healing |
| Gtf2i | general transcription factor II I | negative regulation of angiogenesis |
| Hdac5 | histone deacetylase 5 | negative regulation of cell migration involved in sprouting angiogenesis |
| Hdac7 | histone deacetylase 7 | positive regulation of cell migration involved in sprouting angiogenesis |

| | | |
|---|---|---|
| Hdac9 | histone deacetylase 9 | positive regulation of cell migration involved in sprouting angiogenesis |
| Hhex | hematopoietically expressed homeobox | negative regulation of angiogenesis |
| Hif1a | hypoxia inducible factor 1 alpha subunit | positive regulation of angiogenesis |
| Hipk2 | homeodomain interacting protein kinase 2 | positive regulation of angiogenesis |
| Hmox1 | heme oxygenase 1 | positive regulation of angiogenesis |
| Hpse | heparanase | angiogenesis involved in wound healing |
| Hs6st1 | heparan sulfate 6-O-sulfotransferase 1 | angiogenesis |
| Hspa4 | heat shock protein family A member 4 | positive regulation of angiogenesis |
| Hspb1 | heat shock protein family B (small) member 1 | positive regulation of angiogenesis |
| Hspb6 | heat shock protein family B (small) member 6 | positive regulation of angiogenesis |
| Hyal1 | hyaluronoglucosaminidase 1 | positive regulation of angiogenesis |
| Il15 | interleukin 15 | angiogenesis |
| Il18 | interleukin 18 | angiogenesis |
| Il1a | interleukin 1 alpha | positive regulation of angiogenesis |
| Il1b | interleukin 1 beta | positive regulation of angiogenesis |
| Isl1 | ISL LIM homeobox 1 | positive regulation of angiogenesis |
| Itga5 | integrin subunit alpha 5 | positive regulation of sprouting angiogenesis |
| Itgav | integrin subunit alpha V | angiogenesis |
| Itgb1 | integrin subunit beta 1 | cell migration involved in sprouting angiogenesis |
| Itgb1bp1 | integrin subunit beta 1 binding protein 1 | blood vessel endothelial cell proliferation involved in sprouting angiogenesis |
| Itgb2 | integrin subunit beta 2 | positive regulation of angiogenesis |
| Itgb3 | integrin subunit beta 3 | positive regulation of angiogenesis |
| Jak1 | Janus kinase 1 | positive regulation of sprouting angiogenesis |
| Jam3 | junctional adhesion molecule 3 | angiogenesis |
| Jmjd6 | arginine demethylase and lysine hydroxylase | sprouting angiogenesis |
| Jun | Jun proto-oncogene, AP-1 transcription factor subunit | angiogenesis |
| Kctd10 | potassium channel tetramerization domain containing 10 | angiogenesis |
| Kdr | kinase insert domain receptor | positive regulation of angiogenesis |
| Klf4 | Kruppel like factor 4 | negative regulation of cell migration involved in sprouting angiogenesis |
| Krit1 | KRIT1, ankyrin repeat containing | negative regulation of angiogenesis |
| Lef1 | lymphoid enhancer binding factor 1 | branching involved in blood vessel morphogenesis |
| Lemd3 | LEM domain containing 3 | angiogenesis |
| Lepr | leptin receptor | angiogenesis |
| Lgals3 | galectin 3 | positive regulation of angiogenesis |
| Lif | LIF, interleukin 6 family cytokine | negative regulation of angiogenesis |

| | | |
|---|---|---|
| Lrg1 | leucine-rich alpha-2-glycoprotein 1 | positive regulation of angiogenesis |
| Map2k5 | mitogen activated protein kinase kinase 5 | negative regulation of cell migration involved in sprouting angiogenesis |
| Map3k7 | mitogen activated protein kinase kinase kinase 7 | angiogenesis |
| Mapk14 | mitogen activated protein kinase 14 | angiogenesis |
| Med1 | mediator complex subunit 1 | angiogenesis |
| Meis1 | Meis homeobox 1 | angiogenesis |
| Mfge8 | milk fat globule-EGF factor 8 protein | angiogenesis |
| Mmp14 | matrix metallopeptidase 14 | angiogenesis |
| Mmp2 | matrix metallopeptidase 2 | angiogenesis |
| Mmrn2 | multimerin 2 | negative regulation of cell migration involved in sprouting angiogenesis |
| Mtdh | metadherin | positive regulation of angiogenesis |
| Ncl | nucleolin | angiogenesis |
| Ndnf | neuron-derived neurotrophic factor | angiogenesis |
| Nf1 | neurofibromin 1 | negative regulation of angiogenesis |
| Nfatc3 | nuclear factor of activated T-cells 3 | branching involved in blood vessel morphogenesis |
| Nfatc4 | nuclear factor of activated T-cells 4 | branching involved in blood vessel morphogenesis |
| Nfe2l2 | nuclear factor, erythroid 2-like 2 | positive regulation of angiogenesis |
| Ngfr | nerve growth factor receptor | negative regulation of angiogenesis |
| Nos3 | nitric oxide synthase 3 | positive regulation of angiogenesis |
| Notch1 | notch 1 | negative regulation of cell migration involved in sprouting angiogenesis |
| Notch2 | notch 2 | glomerular capillary formation |
| Notch3 | notch 3 | glomerular capillary formation |
| Notch4 | notch 4 | positive regulation of angiogenesis |
| Nr4a1 | nuclear receptor subfamily 4, group A, member 1 | cell migration involved in sprouting angiogenesis |
| Nrcam | neuronal cell adhesion molecule | angiogenesis |
| Nrp1 | neuropilin 1 | angiogenesis |
| Nrp2 | neuropilin 2 | angiogenesis |
| Otulin | OTU deubiquitinase with linear linkage specificity | sprouting angiogenesis |
| Parva | parvin, alpha | sprouting angiogenesis |
| Pdcd10 | programmed cell death 10 | negative regulation of cell migration involved in sprouting angiogenesis |
| Pdcd6 | programmed cell death 6 | positive regulation of angiogenesis |
| Pdcl3 | phosducin-like 3 | positive regulation of angiogenesis |
| Pde3b | phosphodiesterase 3B | negative regulation of angiogenesis |
| Pdgfa | platelet derived growth factor subunit A | angiogenesis |

| | | |
|---|---|---|
| Pdgfrb | platelet derived growth factor receptor beta | cell migration involved in coronary angiogenesis |
| Pf4 | platelet factor 4 | negative regulation of angiogenesis |
| Pgk1 | phosphoglycerate kinase 1 | negative regulation of angiogenesis |
| Pik3ca | phosphatidylinositol-4,5-bisphosphate 3-kinase, catalytic subunit alpha | angiogenesis |
| Pik3cb | phosphatidylinositol-4,5-bisphosphate 3-kinase, catalytic subunit beta | angiogenesis involved in wound healing |
| Pik3r6 | phosphoinositide-3-kinase, regulatory subunit 6 | positive regulation of angiogenesis |
| Pknox1 | PBX/knotted 1 homeobox 1 | angiogenesis |
| Plau | plasminogen activator, urokinase | angiogenesis |
| Plcd1 | phospholipase C, delta 1 | angiogenesis |
| Plcd3 | phospholipase C, delta 3 | angiogenesis |
| Plcg1 | phospholipase C, gamma 1 | positive regulation of angiogenesis |
| Plxdc1 | plexin domain containing 1 | angiogenesis |
| Plxnd1 | plexin D1 | angiogenesis |
| Pml | promyelocytic leukemia | negative regulation of angiogenesis |
| Pnpla6 | patatin-like phospholipase domain containing 6 | angiogenesis |
| Pofut1 | protein O-fucosyltransferase 1 | angiogenesis |
| Ppp1r16b | protein phosphatase 1, regulatory subunit 16B | positive regulation of blood vessel endothelial cell proliferation involved in sprouting angiogenesis |
| Ppp3r1 | protein phosphatase 3, regulatory subunit B, alpha | branching involved in blood vessel morphogenesis |
| Prcp | prolylcarboxypeptidase | angiogenesis involved in wound healing |
| Prkca | protein kinase C, alpha | positive regulation of angiogenesis |
| Prkcb | protein kinase C, beta | positive regulation of angiogenesis |
| Prkd2 | protein kinase D2 | positive regulation of angiogenesis |
| Prkx | protein kinase, X-linked | angiogenesis |
| Pten | phosphatase and tensin homolog | angiogenesis |
| Ptgs2 | prostaglandin-endoperoxide synthase 2 | angiogenesis |
| Ptk2 | protein tyrosine kinase 2 | angiogenesis |
| Ptk2b | protein tyrosine kinase 2 beta | positive regulation of angiogenesis |
| Ptn | pleiotrophin | negative regulation of angiogenesis |
| Ptprm | protein tyrosine phosphatase, receptor type, M | negative regulation of angiogenesis |
| Ramp1 | receptor activity modifying protein 1 | angiogenesis |
| Ramp2 | receptor activity modifying protein 2 | angiogenesis |
| Rapgef3 | Rap guanine nucleotide exchange factor 3 | positive regulation of angiogenesis |
| Rasip1 | Ras interacting protein 1 | angiogenesis |
| Rbm15 | RNA binding motif protein 15 | branching involved in blood vessel morphogenesis |

| | | |
|---|---|---|
| Rbpj | recombination signal binding protein for immunoglobulin kappa J region | angiogenesis |
| Rhoa | ras homolog family member A | negative regulation of cell migration involved in sprouting angiogenesis |
| Rhob | ras homolog family member B | positive regulation of angiogenesis |
| Rnf213 | ring finger protein 213 | angiogenesis |
| Rock1 | Rho-associated coiled-coil containing protein kinase 1 | negative regulation of angiogenesis |
| Rock2 | Rho-associated coiled-coil containing protein kinase 2 | negative regulation of angiogenesis |
| Rora | RAR-related orphan receptor A | angiogenesis |
| Rras | RAS related | positive regulation of angiogenesis |
| Rtn4 | reticulon 4 | angiogenesis |
| Runx1 | runt-related transcription factor 1 | positive regulation of angiogenesis |
| S1pr1 | sphingosine-1-phosphate receptor 1 | angiogenesis |
| Sars | seryl-tRNA synthetase | negative regulation of angiogenesis |
| Sash1 | SAM and SH3 domain containing 1 | positive regulation of angiogenesis |
| Sat1 | spermidine/spermine N1-acetyl transferase 1 | angiogenesis |
| Scg2 | secretogranin II | angiogenesis |
| Sema3e | semaphorin 3E | negative regulation of angiogenesis |
| Sema4a | semaphorin 4A | negative regulation of angiogenesis |
| Sema5a | semaphorin 5A | positive regulation of angiogenesis |
| Serpine1 | serpin family E member 1 | angiogenesis |
| Setd2 | SET domain containing 2 | angiogenesis |
| Sfrp2 | secreted frizzled-related protein 2 | positive regulation of angiogenesis |
| Sh2b3 | SH2B adaptor protein 3 | negative regulation of sprouting angiogenesis |
| Shc1 | SHC adaptor protein 1 | angiogenesis |
| Sirt1 | sirtuin 1 | positive regulation of angiogenesis |
| Sirt6 | sirtuin 6 | positive regulation blood vessel branching |
| Slit2 | slit guidance ligand 2 | cell migration involved in sprouting angiogenesis |
| Sox17 | SRY box 17 | angiogenesis |
| Sox18 | SRY box 18 | angiogenesis |
| Sparc | secreted protein acidic and cysteine rich | negative regulation of angiogenesis |
| Spred1 | sprouty-related, EVH1 domain containing 1 | negative regulation of cell migration involved in sprouting angiogenesis |
| Srf | serum response factor | cell migration involved in sprouting angiogenesis |
| Srpk2 | SRSF protein kinase 2 | angiogenesis |
| Stab1 | stabilin 1 | negative regulation of angiogenesis |
| Stard13 | StAR-related lipid transfer domain containing 13 | negative regulation of sprouting angiogenesis |
| Stat1 | signal transducer and activator of transcription 1 | negative regulation of angiogenesis |

| | | |
|---|---|---|
| Stk4 | serine/threonine kinase 4 | branching involved in blood vessel morphogenesis |
| Sulf1 | sulfatase 1 | negative regulation of angiogenesis |
| Syk | spleen associated tyrosine kinase | angiogenesis |
| Synj2bp | synaptojanin 2 binding protein | negative regulation of angiogenesis |
| Tal1 | TAL bHLH transcription factor 1, erythroid differentiation factor | angiogenesis |
| Tcf4 | transcription factor 4 | negative regulation of angiogenesis |
| Tek | TEK receptor tyrosine kinase | angiogenesis |
| Tert | telomerase reverse transcriptase | positive regulation of angiogenesis |
| Tgfbi | transforming growth factor, beta induced | angiogenesis |
| Tgfbr1 | transforming growth factor, beta receptor 1 | angiogenesis |
| Tgfbr2 | transforming growth factor, beta receptor 2 | positive regulation of angiogenesis |
| Thy1 | Thy-1 cell surface antigen | angiogenesis |
| Tie1 | tyrosine kinase with immunoglobulin-like and EGF-like domains 1 | negative regulation of angiogenesis |
| Tnfrsf1a | TNF receptor superfamily member 1A | positive regulation of angiogenesis |
| Tspan12 | tetraspanin 12 | angiogenesis |
| Ubp1 | upstream binding protein 1 | angiogenesis |
| Vash1 | vasohibin 1 | negative regulation of angiogenesis |
| Vegfa | vascular endothelial growth factor A | positive regulation of angiogenesis |
| Vegfb | vascular endothelial growth factor B | positive regulation of angiogenesis |
| Vezf1 | vascular endothelial zinc finger 1 | angiogenesis |
| Vhl | von Hippel-Lindau tumor suppressor | angiogenesis |
| Wars | tryptophanyl-tRNA synthetase | angiogenesis |
| Wasf2 | WAS protein family, member 2 | angiogenesis |
| Xbp1 | X-box binding protein 1 | positive regulation of vascular wound healing |
| Zc3h12a | zinc finger CCCH type containing 12A | positive regulation of angiogenesis |

**Table S3: Microglial genes involved in dilation of the vasculature.**

Genes expressed by retinal microglia were compared against genes involved in the regulation of blood vessel diameter (GO:0097746, filtered for Rattus norvegicus). Of the 310 rat genes probed, 41 microglial genes were identified as having a role in the dilation of vasculature.

| **Gene ID** | **Gene name** | **Vasoactive action** |
|---|---|---|
| Adcy6 | adenylate cyclase 6 | regulation of blood vessel diameter |
| Adm | adrenomedullin | positive regulation of blood vessel diameter |
| Adora2a | adenosine A2a receptor | vasodilation |
| Adrb1 | adrenoceptor beta 1 | norepinephrine-epinephrine-mediated vasodilation |
| Adrb2 | adrenoceptor beta 2 | norepinephrine-epinephrine-mediated vasodilation |
| Ahr | aryl hydrocarbon receptor | negative regulation of vasoconstriction |
| Alox12 | arachidonate 12-lipoxygenase, 12S type | positive regulation of blood vessel diameter |
| Apoe | apolipoprotein E | vasodilation |
| Atg5 | autophagy related 5 | vasodilation |
| Bbs2 | Bardet-Biedl syndrome 2 | vasodilation |
| Bmpr2 | bone morphogenetic protein receptor type 2 | negative regulation of vasoconstriction |
| Cx3cl1 | C-X3-C motif chemokine ligand 1 | negative regulation of vasoconstriction |
| Dock4 | dedicator of cytokinesis 4 | negative regulation of vascular smooth muscle contraction |
| Drd1 | dopamine receptor D1 | vasodilation |
| Gch1 | GTP cyclohydrolase 1 | vasodilation |
| Gpx1 | glutathione peroxidase 1 | vasodilation |
| Gucy1a3 | guanylate cyclase 1 soluble subunit alpha 3 | relaxation of vascular smooth muscle |
| Hbegf | Heparin binding EGF-like growth factor | vasodilation |
| Hif1a | hypoxia inducible factor 1 alpha subunit | negative regulation of vasoconstriction |
| Hmgcr | 3-hydroxy-3-methylglutaryl-CoA reductase | negative regulation of blood vessel diameter |
| Hspa1b | heat shock protein family A (Hsp70) member 1B | negative regulation of vasoconstriction |
| Kat2b | lysine acetyltransferase 2B | positive regulation of blood vessel diameter |
| Kcnj8 | potassium voltage-gated channel subfamily J member 8 | vasodilation |
| Kcnma1 | potassium calcium-activated channel subfamily M alpha 1 | vasodilation |
| Kdr | kinase insert domain receptor | positive regulation of blood vessel diameter |
| Map2k1 | mitogen activated protein kinase kinase 1 | vasodilation |
| Mmp2 | matrix metallopeptidase 2 | negative regulation of vasoconstriction |
| Mrvi1 | murine retrovirus integration site 1 homolog | relaxation of vascular smooth muscle |
| P2rx1 | purinergic receptor P2X 1 | regulation of vascular smooth muscle contraction |
| P2rx4 | purinergic receptor P2X 4 | vasodilation |
| P2ry1 | purinergic receptor P2Y1 | regulation of blood vessel diameter |
| P2ry2 | purinergic receptor P2Y2 | regulation of blood vessel diameter |

| | | |
|---|---|---|
| Pla2g6 | phospholipase A2 group VI | positive regulation of blood vessel diameter |
| Plod3 | procollagen-lysine, 2-oxoglutarate 5-dioxygenase 3 | vasodilation |
| Ppard | peroxisome proliferator-activated receptor delta | positive regulation of blood vessel diameter |
| Ptk2 | protein tyrosine kinase 2 | positive regulation of blood vessel diameter |
| Ptprm | protein tyrosine phosphatase, receptor type, M | positive regulation of blood vessel diameter |
| Scpep1 | serine carboxypeptidase 1 | positive regulation of blood vessel diameter |
| Sirt1 | sirtuin 1 | positive regulation of blood vessel diameter |
| Sod1 | superoxide dismutase 1 | vasodilation |
| Sod2 | superoxide dismutase 2 | acetylcholine-mediated vasodilation |

**Table S4: Microglial genes involved in the constriction of vasculature.**
Genes expressed by retinal microglia were compared against genes involved in the regulation of blood vessel diameter (GO:0097746, filtered for Rattus norvegicus). Of the 310 rat genes probed, 39 microglial genes were identified as having a role in the constriction of vasculature.

| Gene ID | Gene name | Vasoactive action |
|---|---|---|
| Abl1 | ABL proto-oncogene 1, non-receptor tyrosine kinase | positive regulation of vasoconstriction |
| Adora1 | adenosine A1 receptor | negative regulation of blood vessel diameter |
| Agt | angiotensinogen | angiotensin-mediated vasoconstriction |
| Akt1 | AKT serine/threonine kinase 1 | positive regulation of vasoconstriction |
| Alox5 | arachidonate 5-lipoxygenase | positive regulation of vasoconstriction |
| Asic2 | acid sensing ion channel subunit 2 | regulation of vasoconstriction |
| Atp1a2 | ATPase Na+/K+ transporting subunit alpha 2 | regulation of vasoconstriction |
| Avpr1a | arginine vasopressin receptor 1A | positive regulation of vasoconstriction |
| Cacna1g | calcium voltage-gated channel subunit alpha1 G | artery smooth muscle contraction |
| Cav1 | caveolin 1 | Positive regulation of vasoconstriction |
| Cd38 | CD38 molecule | positive regulation of vasoconstriction |
| Chga | chromogranin A | negative regulation of blood vessel diameter |
| Cysltr1 | cysteinyl leukotriene receptor 1 | positive regulation of vasoconstriction |
| Ece1 | endothelin converting enzyme 1 | positive regulation of vasoconstriction |
| Edn1 | endothelin 1 | positive regulation of vasoconstriction |
| Edn3 | endothelin 3 | positive regulation of vasoconstriction |
| Ednrb | endothelin receptor type B | positive regulation of vasoconstriction |
| F2r | coagulation factor II (thrombin) receptor | positive regulation of vasoconstriction |
| Icam1 | intercellular adhesion molecule 1 | positive regulation of vasoconstriction |
| Kcna5 | potassium voltage-gated channel subfamily A member 5 | regulation of vasoconstriction |
| Manf | mesencephalic astrocyte-derived neurotrophic factor | vasoconstriction of artery |
| Mkks | McKusick-Kaufman syndrome | artery smooth muscle contraction |
| Nos3 | nitric oxide synthase 3 | positive regulation of vasoconstriction |
| Per2 | period circadian clock 2 | regulation of vasoconstriction |
| Prkcq | protein kinase C, theta | regulation of vasoconstriction |
| Ptafr | platelet-activating factor receptor | positive regulation of vasoconstriction |
| Ptgs1 | prostaglandin-endoperoxide synthase 1 | positive regulation of vasoconstriction |
| Ptgs2 | prostaglandin-endoperoxide synthase 2 | positive regulation of vasoconstriction |
| Rap1gds1 | Rap1 GTPase-GDP dissociation stimulator 1 | vascular smooth muscle contraction |
| Rgs2 | regulator of G-protein signaling 2 | positive regulation of vasoconstriction |
| Rhoa | ras homolog family member A | angiotensin-mediated vasoconstriction |
| Serpinf2 | serpin family F member 2 | regulation of blood vessel diameter by renin-angiotensin |
| Shc1 | SHC adaptor protein 1 | positive regulation of vasoconstriction |

| | | |
|---|---|---|
| Slc8a1 | solute carrier family 8 member A1 | vascular smooth muscle contraction |
| Smpd3 | sphingomyelin phosphodiesterase 3 | artery smooth muscle contraction |
| Snta1 | syntrophin, alpha 1 | regulation of vasoconstriction by circulating norepinephrine |
| Tbxas1 | thromboxane A synthase 1 | positive regulation of vasoconstriction |
| Trpm4 | transient receptor potential cation channel, subfamily M, member 4 | vasoconstriction |
| Wdr35 | WD repeat domain 35 | negative regulation of blood vessel diameter |

**Table S5. Weight and blood glucose levels for control and STZ-treated rats over 4 weeks.** Blood glucose and animal weight was determined biweekly and data are expressed as mean values ± standard deviation, $n = 28$, *$p < 0.0001$ STZ compared to control, †$p < 0.01$ within treatment group temporal analysis.

| | **Week** | **0** | **2** | **4** |
|---|---|---|---|---|
| **Weight (g)** | Control | 169.7 ± 24.9 | 209.9 ± 15.8† | 227.7 ± 21.2† |
| | STZ | 168.2 ± 19.8 | 181.7 ± 22.1* | 183.4 ± 29.8* |
| **Blood glucose (mmol/L)** | Control | 7.8 ± 0.7 | 7.0 ± 0.6 | 7.2 ± 1.1 |
| | STZ | 22.9 ± 5.5* | 26.5 ± 5.9*† | 31.4 ± 2.0*† |

**Table S6. Microglial genes involved in positive regulation of inflammation**

After 4 weeks diabetes, differentially expressed retinal microglial genes were compared against genes involved in positive regulation of inflammation (GO: 0050729, filtered for Rattus norvegicus). Of the 120 rat genes probed, 15 microglial genes were identified as having a role in inflammation.

| **Gene ID** | **Gene name** | **Fold Change** |
|---|---|---|
| S100a8 | Protein S100-A8 | 19.90 |
| S100a9 | Protein S100-A9 | 13.14 |
| Adam8 | ADAM metallopeptidase domain 8 | 5.23 |
| AC117869 | Lysine--tRNA ligase | 4.25 |
| AC103574 | T cell-interacting,-activating receptor on myeloid cells 1 | 2.12 |
| Ccl6 | C-C motif chemokine 6 | 1.95 |
| Gpsm3 | G-protein-signaling modulator 3 | 1.88 |
| Vamp8 | Vesicle-associated membrane protein 8 | 1.80 |
| Alox5ap | Arachidonate 5-lipoxygenase-activating protein | 1.56 |
| Tgm2 | Tissue-type transglutaminase | -7.49 |
| Bst1 | ADP-ribosyl cyclase/cyclic ADP-ribose hydrolase 2 | 7.87 |
| Agt | Angiotensinogen | 2.40 |
| Anxa1 | Annexin A1 | 1.89 |
| Tmsb4x | Thymosin beta-4 | 1.85 |
| Ndufc2 | NADH dehydrogenase [ubiquinone] 1 subunit C2 | 1.64 |

**Table S7. Microglial genes involved in negative regulation of inflammation**

After 4 weeks diabetes, differentially expressed retinal microglial genes were compared against genes involved in negative regulation of inflammation (GO:0050728, filtered for Rattus norvegicus). Of the 139 rat genes probed, 12 microglial genes were identified as being negative regulators of inflammation.

| **Gene ID** | **Gene name** | **Fold Change** |
|---|---|---|
| Nlrp12 | NLR family, pyrin domain-containing 12 | 20.22 |
| Slpi | RCG32428, isoform CRA_a | 10.03 |
| Pglyrp1 | Peptidoglycan recognition protein 1 | 5.10 |
| Isl1 | Insulin gene enhancer protein ISL-1 | 3.42 |
| Spn | Leukosialin (Fragment) | 2.26 |
| Metrnl | Meteorin-like protein | 2.23 |
| Siglec8 | RCG54314 | 1.49 |
| Bst1 | ADP-ribosyl cyclase/cyclic ADP-ribose hydrolase 2 | 7.87 |
| Agt | Angiotensinogen | 2.40 |
| Anxa1 | Annexin A1 | 1.89 |
| Tmsb4x | Thymosin beta-4 | 1.85 |
| Ndufc2 | NADH dehydrogenase [ubiquinone] 1 subunit C2 | 1.64 |

**Movie S1 (separate file).**

**Video S1: *Ex vivo* Cx3cr1$^{GFP/+}$ retinal preparation during PBS and fractalkine exposure.**
A retinal *ex vivo* explant was isolated from Cx3cr1$^{GFP/+}$ mice as per materials and methods and exposed to PBS (0-6 minutes), followed by fractalkine (6 - 16 minutes). Vessel diameters were calculated from areas of microglial contact, in addition to those areas that showed no contact (see highlighted regions in Fig. 2*A)*. Quantitative data from *n* = 6 animals is shown in Fig. 2*B*.

**Video S2: Imaris image rendering of microglial-pericyte contact.**
A retinal wholemount from an NG2-DsRed reporter mouse was used to assess microglial-pericyte contact.  and pericytes (DsRed, red), microglia (Iba-1, green), and endothelial cells (CD31, blue) labelled. Where a microglial process was observed to make contact with a pericyte, an Imaris rendering of the area was undertaken showing microglial-pericyte contact.
